# Genomic Changes During the Evolution of the *Coxiella* Genus Along the Parasitism-Mutualism Continuum

**DOI:** 10.1101/2022.10.26.513839

**Authors:** Diego Santos-Garcia, Olivier Morel, Hélène Henri, Adil El Filali, Marie Buysse, Valérie Noël, Karen D. McCoy, Yuval Gottlieb, Lisa Klasson, Lionel Zenner, Olivier Duron, Fabrice Vavre

## Abstract

The Coxiellaceae family is composed of five genera showing lifestyles ranging from free-living to symbiosis. Among them, *Coxiella burnetii* is a well-known pathogen causing Q fever in humans. This bacterium presents both intracellular (parasitic) and environmental (resistant) forms. Recently, several environmental *Coxiella* genomes have been reported, among which several have come from intracellular mutualistic symbionts of ticks, termed *Coxiella*-like endosymbionts. We sequenced two new *Coxiella*-LE genomes from *Dermacentor marginatus* (CLEDm) and *Ornithodoros maritimus* (CLEOmar) ticks, the latter belonging to the *C. burnetii* lineage. Using these newly sequenced *Coxiella*-LEs and 43 Coxiellaceae genomes, we conducted comparative genomic and phylogenomic analyses to increase our knowledge of *C. burnetii* pathogenicity and the emergence of *Coxiella*-LEs. Results highlight the probably parasitic nature of the common ancestor of the Coxiellaceae. Indeed, the virulence factor Dot/Icm T4 Secretion System is present in most, but not all, Coxiellaceae. Whereas it is part of a putative pathogenic island in *C. burnetii*, it has been entirely lost or inactivated in *Coxiella*-LEs, suggesting its importance in pathogenesis. Additionally, we found that a Sha/Mrp antiporter was laterally acquired in the *C. burnetii* lineage. This antiporter might be involved in alkali resistance and the development of the resistant form that is able to persist in the environment for long periods of time. The Sha operon is eroded or absent in *Coxiella*-LEs. Finally, we found that all *Coxiella* representatives produce B vitamins and co-factors indicating a pre-adaptation of *Coxiella* to mutualism with hematophagous arthropods. Accordingly, the ancestor of *C. burnetii* and *Coxiella*-LEs was likely a parasitic bacterium able to manipulate its host cell and to produce vitamins and co-factors for its own use.

## Introduction

Our current view of the Coxiellaceae family (Gammaproteobacteria: Legionellales) is largely limited to *Coxiella burnetii*. This intracellular pathogen of vertebrates, including humans, is the causative agent of Q fever, a worldwide zoonosis of concern to domestic ruminants, which has a high economic burden (Kampschreur et al., 2014; Madariaga et al., 2003). However, recent ecological surveys have highlighted that the Coxiellaceae family is very diverse, with at least five genera mainly composed of bacteria found in aquatic environments or associated with arthropods (Duron, Doublet, et al., 2018). The emerging picture is that members of this family can interact in various ways with animal hosts, ranging from obligatory mutualism with arthropods (Duron and Gottlieb, 2020) to obligate parasitism with vertebrates, as described for *C. burnetii* (Voth and Heinzen, 2007). Other species, such as those of the genera *Aquicella* and *Berkiella*, are associated with amoebae living in aquatic environments (Mehari et al., 2016; Santos et al., 2003), while still others are defensive symbionts, such as *Rickettsiella viridis* in aphids (Łukasik et al., 2013; Tsuchida et al., 2010). Several putative environmental *Coxiella* metagenome-assembled genomes (MAGs) have also been reported from marine and groundwater samples (Anantharaman et al., 2016).

Coxiellaceae showing high homology to *C. burnetii* have been identified within ticks (Acari: Ixodida) and classified as *Coxiella*-like endosymbionts (hereafter *Coxiella*-LE) (Buysse and Duron, 2021; Klyachko et al., 2007; Lalzar, Harrus, et al., 2012; Liu et al., 2013; Mediannikov et al., 2003). *C. burnetii* and all *Coxiella*-LE together form a monophyletic clade separated from other members of the *Coxiellaceae* family (Duron, Noël, et al., 2015; Gottlieb et al., 2015; Smith et al., 2015). Contrary to the pathogenic lifestyle of *C. burnetii, Coxiella*-LEs are obligate nutritional endosymbionts required for the completion of the tick life cycle, supplementing the tick bloodmeal diet with essential B-vitamins and co-factors (Ben-Yosef et al., 2020; Duron and Gottlieb, 2020; Guizzo et al., 2017; Li et al., 2018; Zhong et al., 2007). All known *Coxiella*-LEs are vertically transmitted from tick females to their offspring during egg maturation and are thus naturally present in most tick neonates (Buysse, Plantard, et al., 2019; Duron, Noël, et al., 2015; Klyachko et al., 2007; Lalzar, Friedmann, et al., 2014). As a consequence of their intracellular lifestyle and their vertical transmission, all *Coxiella*-LEs sequenced genomes are reduced (*∼* 0.6*Mb* to *∼*1.7 Mb) when compared to *C. burnetii* (*∼* 2.0*Mb*). Contrasting genomic variation between *C. burnetii* and *Coxiella*-LEs can therefore enable us to investigate the evolution of host-associated bacteria along the parasitism-mutualism continuum and the mechanisms underlying pathogenicity of *C. burnetii*.

The infectious risk and pathogenicity of *C. burnetii* depends on key biological functions, including the production of an environmentally-resistant morphotype, the ability to manipulate the host cell, and the ability to survive in phagosomes within acidic microenvironments (Schaik et al., 2013). *C. burnetii* indeed presents a biphasic life cycle, where each phase is characterized by a specialized morphotype (Coleman, Fischer, Howe, et al., 2004; Coleman, Fischer, Cockrell, et al., 2007; Minnick and Raghavan, 2012; Voth and Heinzen, 2007). The Small Cell Variant (hereafter SCV) morphotype can resist extreme pressure, temperature, hydric and osmotic stress, UV radiation, and even disinfectants. This morphotype can be considered an endospore because it presents a complex intracellular membrane system, a condensed nucleoid, and a dormant metabolism. Because of these traits, the SCV can persist for long periods in the environment and then infect hosts by inhalation (Coleman, Fischer, Howe, et al., 2004; Coleman, Fischer, Cockrell, et al., 2007). The other morphotype, the Large Cell Variant (LCV), presents the common structure and metabolism of a gram-negative bacterium. In contrast to the SCV, the LCV is sensitive to physical and chemical stress (Minnick and Raghavan, 2012). Due to their resistance, SCVs are considered the primary infective cells, while LCVs correspond to the replicative forms. Indeed, after SCVs are internalized, they fuse to lysosomal vesicles and start to acidify, forming the *Coxiella*-Containing Vacuoles (CCVs). While intracellular pathogens generally hijack the phagocytosis defense system to avoid acidification of the endosome, *C. burnetii* is an acidophilic bacterium able to exploit its host’s phagolysosome. Indeed, *C. burnetii* maintains higher cytoplasmic pH than the phagolysosome: between 5.1, when the external pH is around 2, and 6.9, when the external pH is almost neutral (Hackstadt, 1983). To achieve this, *C. burnetti* uses both active (e.g. acid-resistance systems) and passive (e.g. proteomes enriched in basic residues) mechanisms in order to avoid protein denaturalization (Baker-Austin and Dopson, 2007; Krulwich et al., 2011; Lund et al., 2014). When the pH of the CCV drops to *∼* 4.5, SCVs start to switch to LCVs and replicate, occupying most of the host cell space and depleting all nutrients. New SCVs forms are then produced, the host cell is lysed and the released SCVs infect new cells or host fluids, facilitating the spread of *C. burnetii* (Minnick and Raghavan, 2012; Schaik et al., 2013). Importantly, host cell manipulation by *C. burnetii* depends on a Dot/Icm type IV Secretion System (SS), also present in other Legionellales pathogens such as *Legionella spp*., to translocate different effectors outside the *Coxiella*-Containing Vacuoles (CCV) and inhibit host cell apoptosis (Minnick and Raghavan, 2012; Voth and Heinzen, 2007).

Unlike *C. burnetii, Coxiella*-LEs do not form resistant forms. Moreover, they cannot replicate in vertebrate host cells, nor in acidic axenic media, suggesting they are unable to colonize acidic cell environments (Duron, Noël, et al., 2015). The identification of the genomic bases of these differences and the evolutionary origin of the functions required for *C. burnetii* pathogenesis may help us understand the specific biology of the pathogen and the evolutionary transition that occurred during the evolution of the Coxiellaceae family. Recently, a *Coxiella*-LE from the soft tick *Ornithodoros amblus*, which is closely related to *C. burnetii*, was sequenced (Brenner et al., 2021) The analysis of this genome highlighted different features, and notably the presence of an inactive Dot/Icm T4SS, also detected in some other *Coxiella*-LEs (Buysse and Duron, 2021; Gottlieb et al., 2015). This suggests that *Coxiella*-LEs derive from pathogenic ancestors (Brenner et al., 2021).

To test this hypothesis and better understand evolution within the Coxiellaceae, we sequenced two novel *Coxiella*-LE genomes from two tick species, the first associated with the soft tick *O. maritimus* and the second with the hard tick *Dermacentor marginatus*. Hard and soft ticks refer to the two major tick families (Ixodidae and Argasidae, respectively). *Coxiella*-LEs associated with these two families have different evolutionary histories (Brenner et al., 2021; Duron, Binetruy, et al., 2017; Duron, Sidi-Boumedine, et al., 2015). While *Coxiella*-LE from *D. marginatus* is closely related to other *Coxiella*-LEs from hard ticks, the *Coxiella*-LE from *O. maritimus* is included in the lineage of *C. burnetii*. These newly sequenced genomes were compared with other available Coxiellaceae genomes, including other arthropod symbionts (*Rickettsiella*), a human pathogen (*Diplorickettsia*), and different environmental (aquatic) bacteria (*Aquicella, Berkiella*, and several *Coxiella* MAGs).

## Material and methods

### *Coxiella* DNA Enrichment and Sequencing

*Dermacentor marginatus* adult ticks were collected by flagging the vegetation in fields near Poleymieux, France (GPS location: 45.866312, 4.803687). *Ornithodoros maritimus* specimens were sampled in bird nests on Carteau islet, France (GPS location: 43.377769, 4.857693). Both tick species were kept alive at 20^*o*^C and 80% humidity until use. Genomic DNA extractions enriched in *Coxiella*-LE DNA were obtained as previously described (Duron, Morel, et al., 2018; Gottlieb et al., 2015). Briefly, Malpighian tubules and ovaries were dissected from 10 adult ticks of each species, pooled, and then homogenized in 100*µ*l sterile double-distilled water. The obtained homogenate was diluted and incubated for 1h at 20^*o*^C in 10ml sterile double-distilled water. To remove host nuclei and other cell debris, the homogenate was filtered using a 5*µ*m Minisart filter (Sartorius). The remaining cells in the homogenate were pelleted by centrifugation (15min at 20,000 x g at 4^*o*^C). Total genomic DNA (gDNA) was extracted from the obtained pellet using the DNeasy Blood and Tissue Kit (Qiagen). The obtained gDNA was quantified on Qubit using the dsDNA high-sensitivity kit (Invitrogen). *Coxiella*-LE DNA enriched samples were sequenced using HiSeq2000 technology by Genotoul DNA Services Facility (Castanet-Tolosan, France) using the TruSeq Nano DNA library construction and HiSeq SBS v3 kits (Illumina). For each sample, a total of *∼*15 Gb of 2×100 bp paired-ended sequences were obtained.

### Assembly and Annotation

The Illumina reads were quality screened and trimmed using UrQt v1.0.18 (Modolo and Lerat, 2015). Cleaned reads were assembled into contigs with SPAdes v1.12 (Bankevich et al., 2012) to create a draft genome sequence. Obtained contigs were collapsed with SiLiX v1.2.11 at 95% nucleotide identity (Miele et al., 2011). Bandage v0.8.1 was used to visualize the SPAdes graph assembly and discard contigs from bacteria other than *Coxiella* and to identify repeated regions (Wick et al., 2015). The *Coxiella*-LE genome of *O. maritimus* was left at the draft status because large amounts of repetitive regions were present. For the genome of *Coxiella*LE from *D. marginatus*, PCR-gap closing was performed as previously described Gottlieb et al. (2015).

Genome annotation of *Coxiella*-LE from *O. maritimus* (named strain CLEOmar) and *D. marginatus* (strain CLEDm) was performed by running a DIYA v1.0 custom pipeline (Stewart et al., 2009), as described in Ellegaard et al. (2013). Briefly, the DIYA pipeline included an initial gene calling step using Prodigal (Hyatt et al., 2010), followed by tRNA and rRNA prediction using tRNAscan-SE (Lowe and Eddy, 1997) and RNAmmer (Lagesen et al., 2007), respectively. Pseudogene prediction was performed by GenePrimp (Pati et al., 2010). Potential functions of predicted protein-encoding genes were assigned using BLASTp (Camacho et al., 2009) against the Uniprot database (The UniProt Consortium, 2012) and PfamScan with the PFAM database (Punta et al., 2012). Manual curation was conducted using Artemis (Rutherford et al., 2000).

Insertion Sequences (hereafter IS) were predicted with ISsaga (Varani et al., 2011). CLEOmar IS copy numbers were estimated by mapping Illumina reads with Bowtie2 v2.4.2 (--very-sensitive-local preset) (Langmead and Salzberg, 2012) against a database containing a reference copy for each IS and five single copy housekeeping genes (Table S2). Coverage and associated descriptive statistics were calculated with Qualimap v2.2.1 (Okonechnikov et al., 2015). The relative copy numbers of IS elements were obtained using the average coverage of housekeeping genes as a reference.

BUSCO v4.0.6 and the corresponding legionellales_odb10 database (creation date 24-04-2019) were used to assess genome completeness (Seppey et al., 2019). The complete genome of *Coxiella* sp. CLEDm and the draft genome of CLEOmar were deposited at the European Nucleotide Archive (ENA) under accession numbers GCA_907164955 and GCA_907164965, respectively.

### Comparative Genomics, Clusters of Orthologous Proteins Inference, and Phylogenomic Reconstruction

The general functions of proteomes were assigned using BLASTp against the Clusters of Orthologous Groups (COG) database (Tatusov et al., 2003). The metabolic potential was assessed by using the proteomes as input for KAAS (Moriya et al., 2007). Homology between CLEOmar pseudogenes and *C. burnetii* RSA 493 genes was assessed by a reciprocal best hit search strategy using MMseq2 (rbh --search-type 3 --max-seqs 100 --max-accept 10) (Steinegger and Söding, 2017).

The core, shared, and specific Clusters of Orthologous Proteins (hereafter COPs) were inferred for 44 Coxiellaceae from all five genera: *Coxiella* (including four *C. burnetii*, nine *Coxiella*-LE, and 23 environmental *Coxiella* MAGs proteomes), *Aquicella* (two), *Berkiella* (two), *Diplorickettsia* (one), and *Rickettsiella* (three) (Table S1). *Legionella pneumophila* str. Philadelphia 1 was included as an outgroup since this bacterium belongs to the Legionellaceae, the sister family to the Coxiellaceae. Annotated Coxiellaceae genomes were downloaded from RefSeq. Gene calling of unannotated *Coxiella* MAGs was performed with Prokka v1.14.5 (--mincontiglen 200 --gram neg) (Seemann, 2014). COPs were inferred with OrthoFinder v2.3.12 (-M msa -T iqtree) (Emms and Kelly, 2019). Obtained COPs table was queried to retrieve specific subsets of COPs and to check for the presence/absence of COPs in different Coxiellaceae. UpSetR v1.4.0 package available in R v3.6.3 (R Core Team, 2020) was used to plot the different COPs intersections between genomes (Conway et al., 2017). Putatively horizontally transferred genes in *C. burnetii* RSA493 were detected with HGTector v2.03b (-m diamond --aln-method fast) (Zhu et al., 2014).

A first species tree was obtained with OrthoFinder. In brief, 348 individually aligned COPs were selected by OrthoFinder to build a concatenated alignment. Then, the species tree of the Coxiellaceae dataset was inferred using the STAG algorithm (Emms and Kelly, 2019). To obtain node support values, a second species tree was computed as follows: (i) positions with more than 50% of the sequences being gaps in the OrthoFinder concatenated alignment (107812 positions) were filtered out with Gblocks v0.91b (21499 selected positions in 143 blocks) (Castresana, 2000); (ii) IQ-TREE v2.0.3 was used to infer the Maximum Likelihood phylogenomic tree using the best suggested evolutionary model (-m MFP) and ultrafast bootstrap (-bb 1000) and SH-like approximate likelihood ratio test (-alrt 1000) (Kalyaanamoorthy et al., 2017; Nguyen et al., 2015).

Single gene phylogenies were obtained by aligning homologous sequences with MAFFT v7.310 (linsi algorithm) (Katoh et al., 2002), computing the Maximum Likelihood tree with IQ-TREE (same options as described above). FigTree v1.4.4 and InkScape v0.92 were used respectively to plot and modify phylogenetic trees to their final version.

A synteny plot of 458 single copy COPs shared between *C. burnetii* RSA493 and *Coxiella* symbionts of *Amblyomma, Dermacentor*, and *Rhipicephalus* tick species was generated with the genoPlotR v0.8.9 R package (Guy et al., 2010). A synteny plot of the Sha/Mrp antiporter and Dot/Icm T4SS region in selected *Coxiella* was produced with genoPlotR.

Synteny between *C. burnetii* strains was computed using OrthoFinder (Emms and Kelly, 2019). IslandViewer 4 database was used to visualize and predict genomic (pathogenic) islands in *Coxiella burnetii* strains (Bertelli et al., 2017). genoPlotR was used to plot synteny and the genomic location of the Sha/Mrp antiporter, the Dot/Icm T4SS region, and IslandViewer 4 results in *C. burnetii* strains. Figures were adjusted with InkScape.

### Isoelectric Point Prediction

To test for proteome-wide adaptation to acid pH, the Isoelectric Points (pI) of all proteins encoded by the different Coxiellaceae were estimated using IPC v1.0 (Kozlowski, 2016). The IPC 2.0 web-server was used to predict pI and charge of glutamate decarboxylase A (GadA) and B (GadB), and Aspartate 1-decarboxylase PanD from all Coxiellaceae, several acidophiles (*Listeria monocytogenes, Lactococcus lactis, Shigella flexneri Mycobacterium tuberculosis*, and *Helicobacter pylori*), and *Escherichia coli* as a neutrophile (Kozlowski, 2021). All statistical tests were performed in R (R Core Team, 2020).

## Results

### *Coxiella spp*. CLEOmar and CLEDm Genomic Features

The genome of *Coxiella*-LE from *D. marginatus* (hereafter CLEDm), was recovered as nine contigs (142X). CLEDm gaps were closed by PCR, resulting in a circular genome of 0.9 Mb with 659 predicted protein-coding genes (CDS), one ribosomal operon, a complete set of tRNAs, 15 putative pseudogenes, and no signal of mobile elements (Table 1).

**Table 1.** General Genomic Features of representative *Coxiella* and *Coxiella*-LE genomes compared to *Coxiella*-LE of *O. maritimus* and *D. marginatus*.Only tick species names are displayed. All tick-hosts belong to the hard ticks family with except *O. maritimus*, a soft tick. ND: No families/copies detected by ISSaga.

| Genome | Strain | Host | Size (Mb) | Genes (CDS) | Pseudogenes | rRNA/tRNA/other RNA | IS families/copies | %GC | Contigs |
| --- | --- | --- | --- | --- | --- | --- | --- | --- | --- |
| *Coxiella*-LE | CLEdm | *Dermacentor marginatus* | 0.9 | 658 | 15 | 3/40/3 | 0/0 | 35.1 | 1 |
| *Coxiella*-LE | CLEOmar | *Ornithodoros maritimus* | 1.83 | 976 | 608 | 3/42/8 | 4/31 | 41.5 | 112 |
| *Coxiella*-LE | AB428 | *Ornithodoros amblyus* | 1.56 | 889 | 660 | 3/42/4 | ND | 40.6 | 101 |
| *C. burnetii* | RSA493 | Mammals | 1.99 | 1833 | 207 | 3/42/NA | 6/32 | 42.6 | 1 |
| *Coxiella*-LE | CRT | *Rhipicephalus turanicus* | 1.73 | 1293 | 337 | 3/47/4 | 0/0 | 38.2 | 1 |
| *Coxiella*-LE | C904 | *Amblyomma americanum* | 0.66 | 565 | 3 | 3/39/2 | 0/0 | 34.6 | 1 |

The genome of *Coxiella*-LE from *O. maritimus* (hereafter CLEOmar) was assembled in 112 contigs (426X average coverage). CLEOmar was left as a draft given the high number of Insertion Sequences (IS), many of which are found at contig edges, and of duplicated regions (Table 1). It contains 976 predicted CDS, 608 pseudogenes, and signatures of active, or recent, IS transposition. A total of four IS families were detected (Table S2). IS1111, from the IS110 subgroup (ssgr), was the most widespread IS with 27 copies. The relative coverage of this IS compared to that of several single-copy genes supports the number of detected copies. The other families presented between one and two copies. However, the number of copies of IS4 ssgr IS10 is underestimated due to its fragmented presence at contig edges. The difficulty in recovering full IS4 ssgr IS10 copies suggests that the identical, or highly similar, copies of this IS are associated with recent transposition events.

Before conducting further analyses, we assessed CLEOmar and CLEDm genome completeness by comparing their BUSCO results to that of selected Coxiellaceae genomes (Table S1, Fig S1). Despite its draft status, the CLEOmar BUSCO score was close to Coxiellaceae genomes of similar size, including *Coxiella*-LE genomes from *Rhipicephalus* tick species. CLEOmar encoded a few more BUSCO genes compared to *Coxiella*-LE AB428 from *O. amblus*. This difference is expected since the latter presents a more reduced genome. Hence, we consider the CLEOmar genome as complete or almost complete.

### Inferring the Coxiellaceae Family Phylogeny

To understand the evolutionary history of the Coxiellaceae family, we first obtained an updated phylogeny of its different members. Phylogenetic relationships between Coxiellaceae were inferred using 348 COPs computed from available Coxiellaceae and environmental relatives (Fig 1). *Aquicella, Rickettsiella*, and *Diplorickettsia* were recovered as a sister clade to *Coxiella. Berkiella* was the most basal Coxiellaceae clade. The phylogeny obtained placed *C. burnetii*, all *Coxiella*-LEs, and the environmental *Coxiella* MAGs as a monophyletic clade with robust node support (100% SH-aLRT and 93% ultrafast bootstrap). Two environmental (groundwater) *Coxiella* MAGs (GCA_001795425 and GCA_001797285) were basal to the subclade containing *C. burnetii* and all *Coxiella*-LEs. Although monophyletic, this subclade was also divided into two groups: one including all *C. burnetii* strains plus *Coxiella*-LEs CLEOmar and AB428 (from *Ornithodoros* soft ticks) and another solely formed by the rest of available *Coxiella*-LEs (from *Amblyomma, Dermacentor*, and *Rhipicephalus* hard ticks).

**Figure 1.**
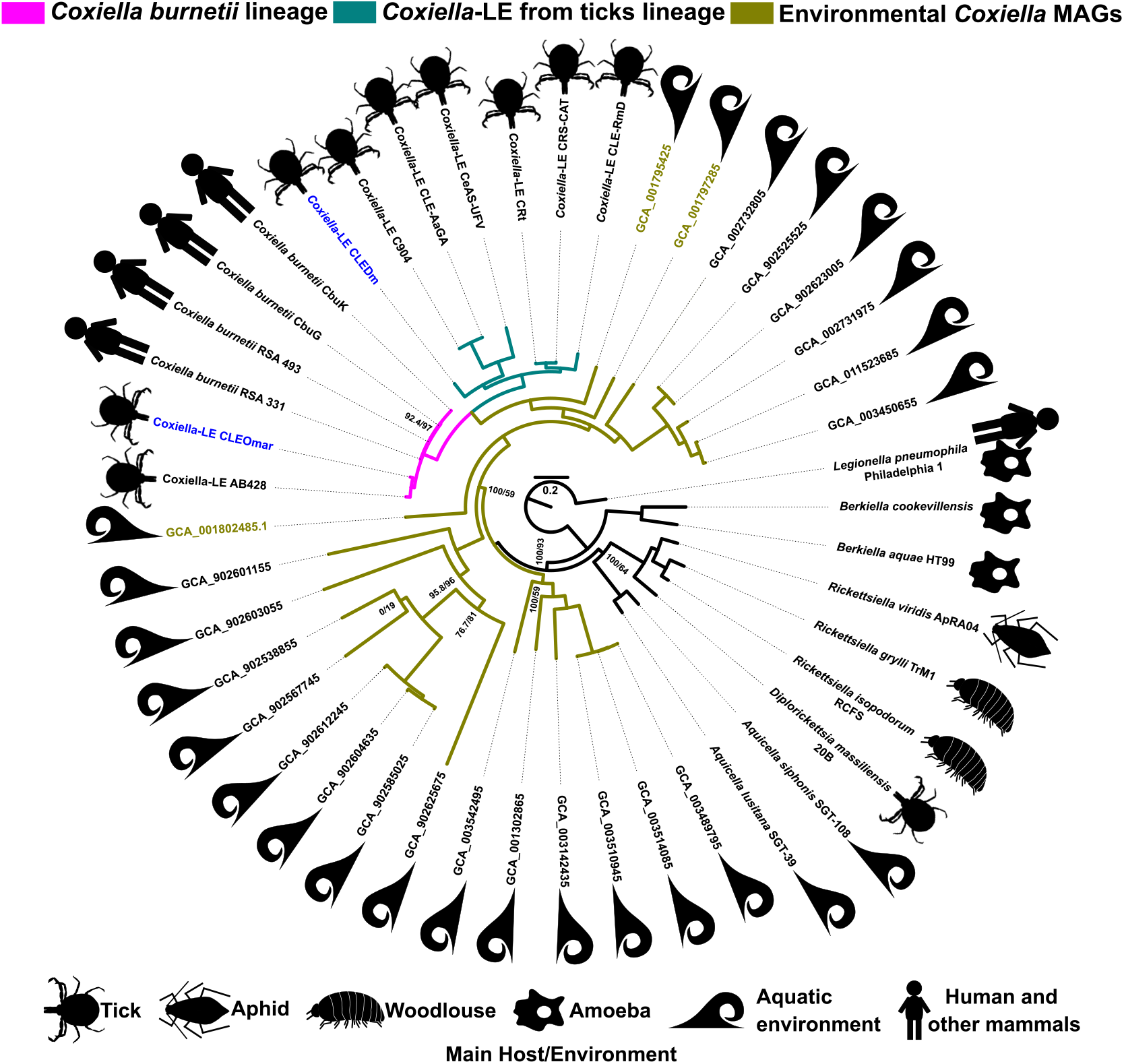
Maximum Likelihood phylogenomic tree of selected Coxiellaceae genomes. The tree was inferred from a concatenated alignment of 348 Clusters of Orthologous Proteins (COPs) under the LG+R6 model. SH-aLRT/ultrafast bootstrap support values numbers are displayed at each node if they are below 99. Newly sequenced *Coxiella*-LE genomes are highlighted in blue. Only those environmental *Coxiella* MAGs highlighted in green were used in further analyses.

### *Coxiella* and *Coxiella*-Like Comparative Genomics

The evolution of gene content between related species with different lifestyles can help us understand transitions between pathogenic and mutualistic relationships. We thus compared the distribution of COPs within the genus *Coxiella* (Table S3). The number of core COPs accounted for up to 21% of the protein-coding genes in *C. burnetii*. This small percentage was expected as the number of core COPs is driven by the most reduced genomes: *Coxiella*-LE strains from *Amblyomma* and CLEDm. The 631 species-specific COPs of *C. burnetii* represented around 34% of its protein-coding genes (Fig 2). The majority of *C. burnetii* specific proteins were assigned to clusters without a defined function (R, S, X) according to COG categories (Fig S2). Out of the 631 species-specific COPs of *C. burnetii*, 300 were identified as pseudogenes in the CLEOmar genome, which indicates their presence in their Most Recent Common Ancestor (MRCA). Genes belonging to R, S, and X COG categories are poorly defined and are generally related to environmental responses. As a large proportion of CLEOmar pseudogenes are in these categories, it seems that these bacteria may be losing their ability to respond to environmental variations. This pattern of genome reduction is similar to the one described in *Coxiella*-LE AB428 (Brenner et al., 2021).

**Figure 2.**
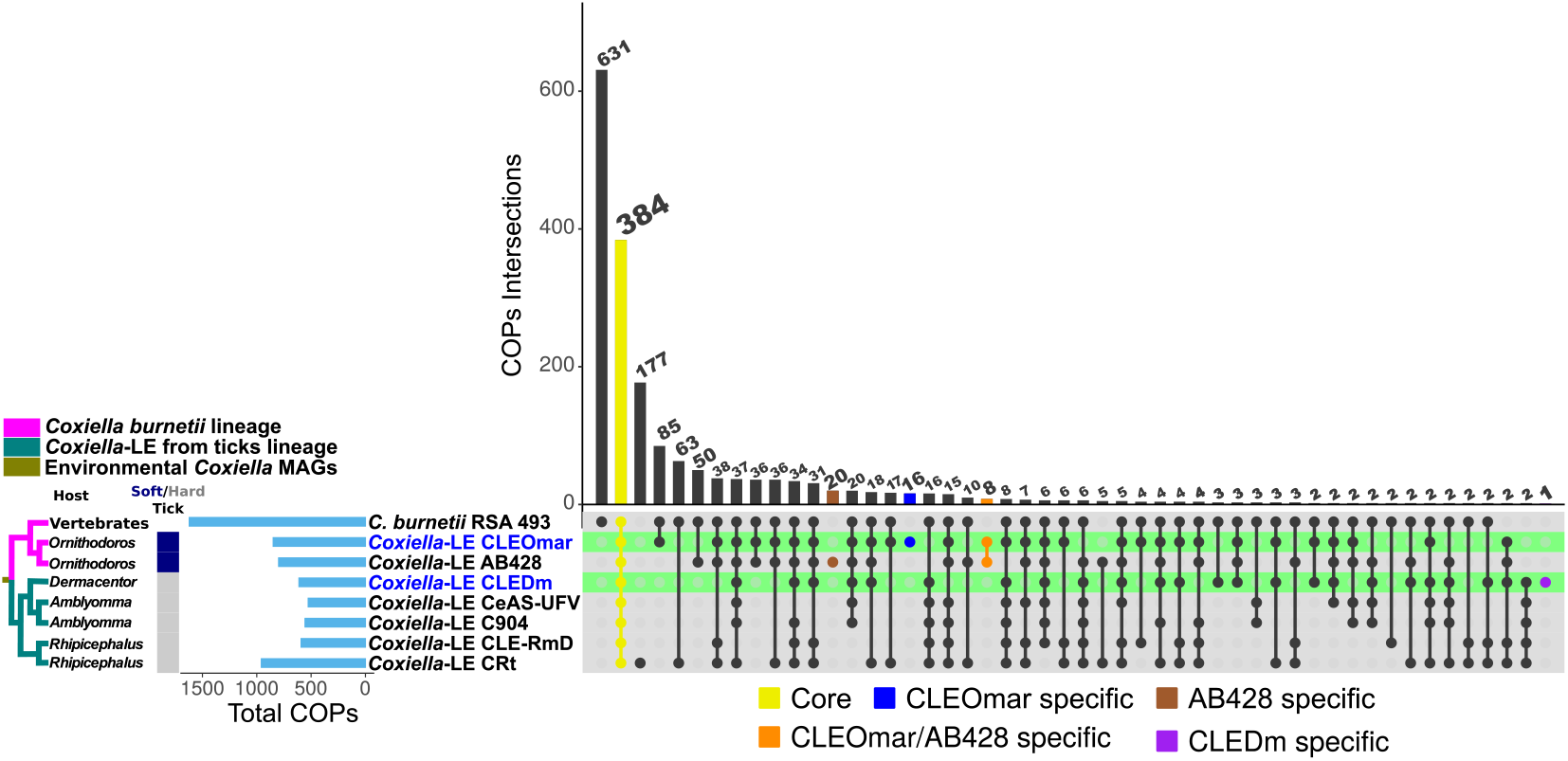
Upset plot displaying shared and specific Clusters of Orthologous Proteins (COPs intersections) between selected *Coxiella* symbionts of ticks and *C. burnetii*. Tick genera and families are displayed on the left. Colored bars denote shared COPs by all included *Coxiella* (yellow), and CLEOmar (blue), AB428 (brown), CLEOmar and AB428 (orange), and CLEDm (purple) specific COPs. Other abbreviations: *C. burnetii* RSA 493, *Coxiella*-LEs AB428 (*Ornithodoros amblus*), C904 (from *A. americanum*), CeAs-UFV (*A. sculptum*), CRt (*Rhipicephalus turanicus*), and CLE-RmD (*R. microplus*). The cladogram on the left represents the phylogenetic relationships of Coxiella species based on Fig 1. Species color coding is as in Fig 1.

In *C. burnetii*, 15 horizontally acquired genes were reported to increase its fitness and virulence (Moses et al., 2017). These genes were assigned to lipopolysaccharide (five genes), fatty acid (seven), biotin (one), and heme (two) biosynthesis pathways (Table S4). As only a few of these are present in Coxiellaceace outside the *Coxiella* clade (*Aquicella lusitana* contains one and *Aq. siphonis* three), most of the genes were probably acquired in the *Coxiella* lineage (Table S4). However, as most were also present in different environmental species of *Coxiella*, they are not specific to *C. burnetii*. Among these genes, only *fabA* and a putative toxin-antitoxin system (CBU_0284-5), likely involved in heme biosynthesis, seem to have been acquired specifically by the *Coxiella*/*Coxiella*-LE lineage (Moses et al., 2017). Several of these horizontally acquired genes are detectable as pseudogenes in reduced *Coxiella*-LE species, suggesting that they are not required for mutualistic relationships.

Core COPs represented more than half of the proteome in highly reduced *Coxiella*-LEs, roughly 58% and 68% in CLEDm and strains from *Amblyomma* tick species, respectively. For larger *Coxiella*-LEs, the percentage was lower but still represented an important part of the proteome: 39% and 30% in CLEOmar and CRt, respectively. Species-specific COPs represented a variable, but relatively small fraction of *Coxiella*-LE proteomes compared to the 34% (631) in *C. burnetii*: 14% (177) in CRt, *∼*2% in CLEOmar (16) and AB428 (20), and *<* 1% in CLEDm (1) and C904 (1). COG classification of the COPs showed that basic cellular processes, such as translation and transcription (J); replication, recombination, and repair (L); or post-translational modifications and chaperonines (O) are retained in reduced *Coxiella*-LE genomes compared to *C. burnetii* (Fig S2). Additionally, co-enzyme transport and metabolism (H) is also retained in reduced *Coxiella*-LE genomes (Fig S2), as already reported in other facultative symbionts suffering a genomic shrinkage and evolving towards a more obligatory status (Manzano-Marín, Lamelas, et al., 2012). For shared COPs, macrosynteny is only conserved between *Coxiella*-LE from *Amblyomma*tick species, except for one re-arrangement detected between strain CoAs and the rest (Fig S3). CLEDm is partially syntenic to *Coxiella*-LE strains from *Amblyomma*. Nonetheless, some conserved regions were detected among *C. burnetii* and all *Coxiella*-LE bacteria.

### Diversity of Coxiellaceae B Vitamin Biosynthesis Potential

The ability to produce B vitamins and co-factors may have played a major in *Coxiella* evolution, especially for the endosymbiotic lineages (Duron and Gottlieb, 2020). The metabolic potential for B vitamin biosynthesis in Coxiellaceae species shows that all are able to produce riboflavin (B2) and lipoic acid. Besides, single-gene phylogenetic trees support the ancestrality of those two pathways despite some incongruencies in the position of different *Coxiella* MAGs (Fig 1, Fig S4, Fig S5, and Supplementary Data). Altogether, our analyses suggest that riboflavin (B2) and lipoic acid production are ancestral traits of the Coxiellaceae family. At the same time, the rest of the B vitamins present a patchy distribution across the family. The most parsimonious explanation for the presence of incomplete pathways (Fig 3) across the different Coxiellaceae clades (Fig 1) is that the Coxiellaceae ancestor was able to produce all B vitamins and co-factors. Then, during evolution, this potential was differentially lost in some genera/lineages (e.g. *Rickettsiella*), but retained in others (*Coxiella*) (Fig 1).

**Figure 3.**
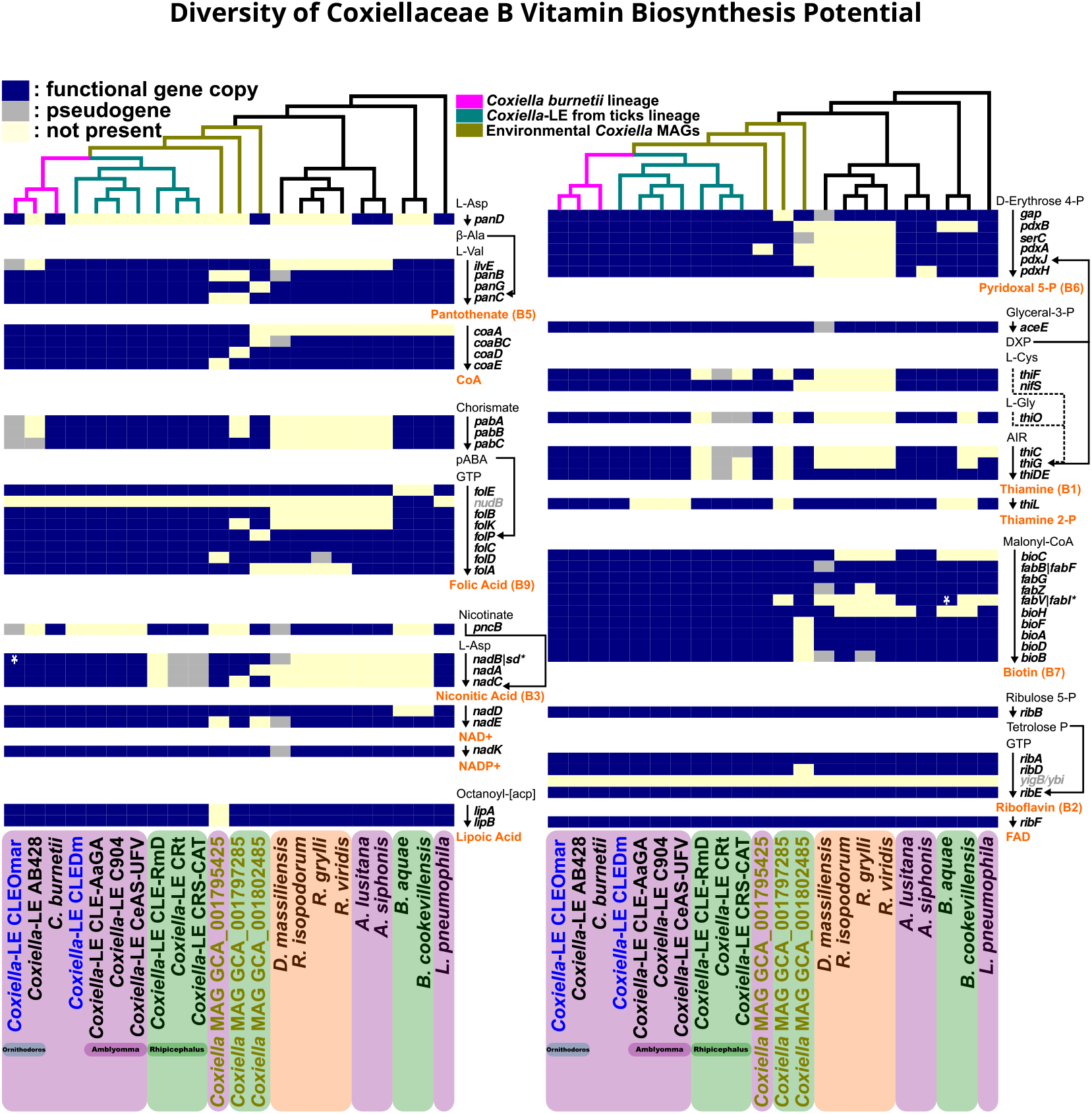
Biosynthetic pathways for B vitamins and co-factors in Coxiellaceae genomes. Three major groups are highlighted according to their metabolic potential: large (purple), medium (green), and reduced (orange). Gene names denoted in gray are rarely found in symbionts providing B vitamins to hematophagous hosts, suggesting unknown alternative enzymatic steps in the pathway. Species color coding is as in Fig 1. Only *C. burnetii* RSA 493 is displayed since all *C. burnetii* strains present the same B vitamin biosynthetic potential. The cladograms on the top represent the phylogenetic relationships of Coxiellaceae species based on Fig 1. Species color coding is as in Fig 1. *: Alternative enzymatic step.

Furthermore, Coxiellaceae can be divided into three major functional groups according to their potential to produce other B vitamins and co-factors (Fig 3 and Table S5). The first functional group includes *C. burnetii*, which presents the largest metabolic potential, together with *Coxiella*-LEs CLEDm, CLEOmar, and AB428, all *Coxiella*-LEs from *Amblyomma* tick species, *Coxiella* MAG GCA_001795425, both *Aquicella* species, and the *L. pneumophila* outgroup (Fig 3). All species in this group can produce almost all B vitamins *de novo* or from intermediate metabolites. While pantothenate (B5), pyridoxine (B6), thiamine (B1), biotin (B7), riboflavin (B2), and lipoic acid pathways are complete in almost all members from this group, nicotinic acid (B3) and folic acid (B9) are only complete in *C. burnetii* and *Coxiella*-LEs (Fig 3). In most single-gene trees from the different biosynthetic pathways (Supplementary Data), *C. burnetii* and *Coxiella*-LEs lineage topology follows the species tree and have *Coxiella* MAGs GCA_001795425, GCA_001797285, and GCA_001802485 as basal clades. Therefore, the *C. burnetii*/*Coxiella*-LEs ancestor encoded the mentioned metabolic pathways. This is even true for the biotin (B7) pathway which is prone to be acquired by HGT (Duron and Gottlieb, 2020). Indeed, single-gene trees suggest that the biotin pathway has been transferred among Coxiellaceae, or even acquired from distant bacterial species. However, the monophyly of biotin genes in the *C. burnetii* and *Coxiella*-LEs lineage support their presence in the last common ancestor of this lineage (Fig S6 and Supplementary Data).

The second and third functional groups present a more restricted metabolic potential. The second group includes *Coxiella*-LEs from *Rhipicephalus* tick species, *Berkiella*, and the two *Coxiella* MAGs GCA_001797285 and GCA_001802485 (Fig 3). The thiamine pathway has been lost in almost all members of this group. In addition, nicotinic acid has been almost lost in *Berkiella*, while *Coxiella*-LEs from *Rhipicephalus* tick species need to import nicotinate to produce NAD^+^/NADP^+^. The biotin pathway seems to be inactive in *Berkiella* and both *Coxiella* MAGs. The third, and last, group includes *Rickettsiella* and *Diplorickettsia* species, which lack the ability to produce thiamine, biotin, folic acid, and pantothenate (Fig 3).

### Evolution of Coxiellaceae Virulence: Phase-Specific proteins and the Dot/Icm System

It is known that *C. burnetii* encodes several proteins which are over- or under-expressed in the different morphotypes (SCV or LCV) and may play important roles in pathogenicity (Coleman, Fischer, Cockrell, et al., 2007). Those proteins are defined, according to their expression profiles in the morphotypes, as LCV^*Hi*^/SCV^*Lo*^ and SCV^*Hi*^/LCV^*Lo*^. Among these phase-specific proteins, the small cell variant protein A (ScvA) and histone-like Hq1 (HcbA) are thought to be involved in nucleoid condensation in SCVs (Coleman, Fischer, Cockrell, et al., 2007). Therefore, we assessed the presence of LCV^*Hi*^/SCV^*Lo*^ and SCV^*Hi*^/LCV^*Lo*^ proteins among the different Coxiellaceae genomes (Table S9). The *scvA* gene was only detected in *C. burnetii*. A functional gene copy of *hcbA* (or *hq1*) was present in CLEOmar, but not in AB428, and a pseudogenized copy was detected in both CRt and CRS-CAT. These two proteins seem to be *C. burnetii*/*Coxiella*-LE clade-specific as they were not found in any other Coxiellaceae analyzed here.

The Dot/Icm has been classified as a Type 4 Secretion System (T4SS) and is essential for the invasion and survival of *C. burnetii* and *Legionella* in their respective hosts. Because of its importance in *C. burnetii* pathogenicity, we investigated the presence of its 25 core proteins in the different Coxiellaceae (Gomez-Valero, Chiner-Oms, et al., 2019). The Dot/Icm T4SS, or traces of it, was detected in almost all Coxiellaceae (Fig 4, Table S6). A few functional genes were detected in *Coxiella*-LEs CRt, CRS-CAT, and CLEOmar. While some pseudogenes were detected in *Coxiella*-LE AB428, no traces of the Dot/Icm T4SS were detected in *Coxiella*-LEs CLEDm and from *Amblyomma* ticks (the most reduced). Nonetheless, the presence of the Dot/Icm T4SS in most Coxiellaceae genomes along all their phylogeny indicates its ancestral state. Although single-gene phylogenetic trees do not completely mimic the species tree, in general their pattern supports the ancestrality of the Dot/Icm T4SS: *Berkiella, Aquicella*, and *Rickettsiella/Diplorickettsia* tend to cluster together in a basal position to a *Coxiella* group, which included most *Coxiella* MAGs and the *C. burnetii*/*Coxiella*-LEs clade (Supplementary Data).

**Figure 4.**
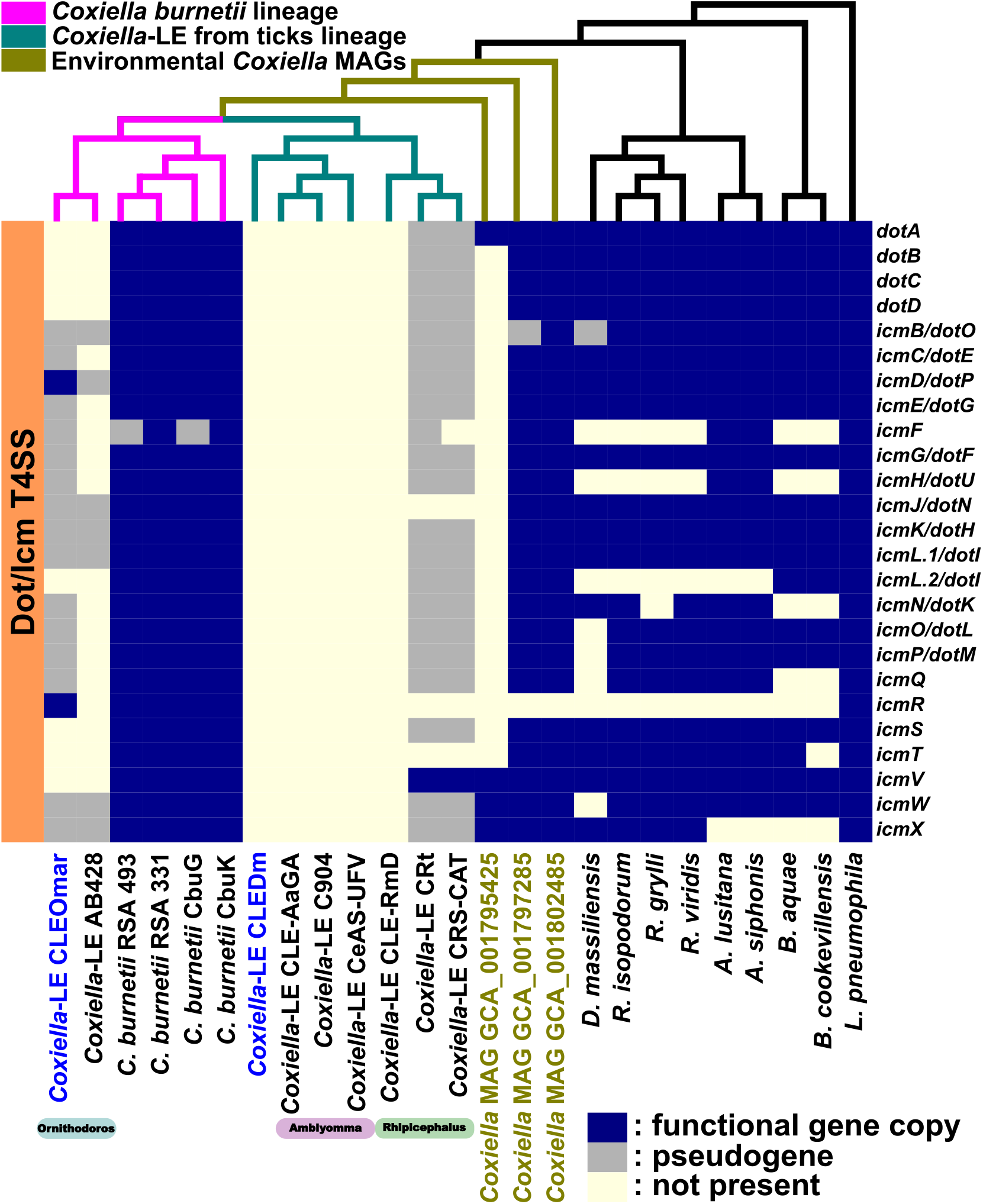
Dot/Icm Type 4 Secretion System presence among selected Coxiellaceae genomes. Only the 25 core components of the Dot/Icm T4SS were included in the analysis (Gomez-Valero, Rusniok, et al., 2019). The cladogram on the top represents the phylogenetic relationships of Coxiellaceae species based on Fig 1. Species color coding is as in Fig 1. Species color coding is as in Fig 1.

In *C. burnetii*, the region containing the Dot/Icm T4SS resembles a pathogenic island (PAI), with the presence of tRNAs, IS elements, direct repeats, horizontally transferred genes (HGT) (Table S7), and virulence factors (Fig 5) (Hacker and Kaper, 2000). Additionally, part of the region is predicted to be a genomic island by IslandViewer 4 in different *C. burnetii* strains (Bertelli et al., 2017) (Table S8, Fig S7). The putative PAI seems to be included in a larger region, around *∼*144 Kb, which has suffered several translocations and inversions in *C. burnetii* strains (Fig S7). Among the predicted HGT (Table S7), it is noticeable the presence of a Sodium Hydrogen/Multiple resistance and pH (Sha/Mrp) antiporter. The Sha/Mrp antiporter is located upstream from the Dot/Icm T4SS and is composed by six genes (*shaABCDEFG*) organized as an operon (Fig 5). This operon might have been acquired from a *Coxiella* relative, such as *Coxiella* sp. GCA_001802485, but its origin is probably from *Legionella* (Fig S8-S14). Indeed, only *B. cookevillensis* encodes another Sha operon (Fig S15), but one which is unrelated to that one of the *C. burnetii* lineage (Fig S8-S14), supporting different HGT events.

**Figure 5.**
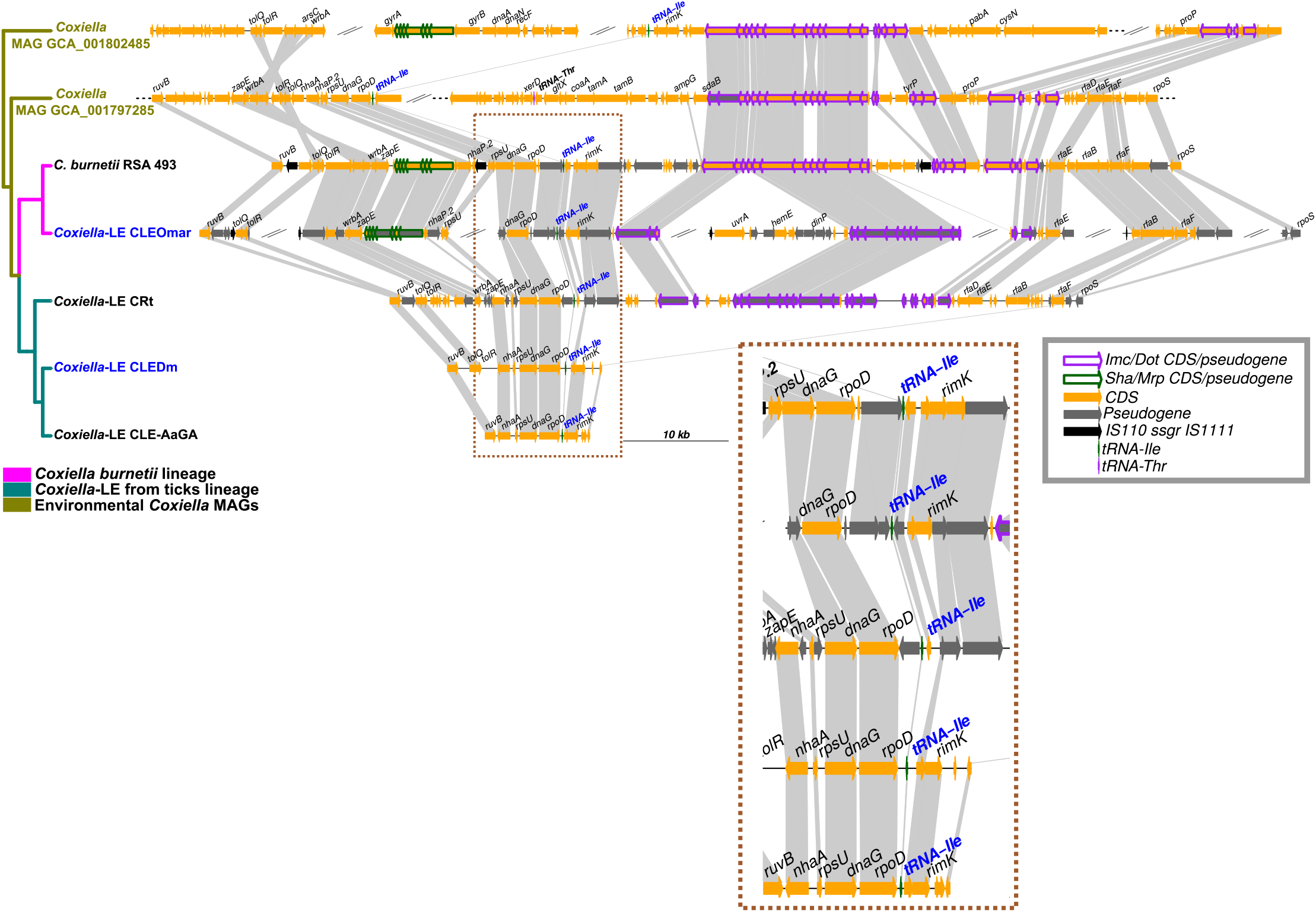
Dot/Icm T4SS genomic region from selected *Coxiella* species compared to *C. burnetii* RSA 493. For draft genomes, only contigs, or regions (denoted as dotted lines) containing Dot/Icm or *Sha* genes are displayed. Different contigs and regions are separated by double slashes. Gray lines connect orthologous genes. Twisted gray lines indicate inversions. Genes annotated as hypothetical proteins or without official names are not displayed. The cladogram on the top represents the phylogenetic relationships of Coxiellaceae species based on Fig 1. Species color coding is as in Fig 1. Species color coding is as in Fig 1.

In general, the putative PAI containing the Dot/Icm T4SS presents the same gene order (microsynteny) in all *C. burnetii* strains, except in RSA 331 (Fig S7). This strain has a small inversion of 6.2 Kb containing six genes, four of them belonging to the Dot/Icm T4SS (*dotA, icmV, icmW, icmX*), and one unrecognizable pseudogene. The inversion is flanked by two identical copies of the same IS, suggesting a relatively recent event. Several regions of the PAI are present in *Coxiella* MAGs GCA_001802485, GCA_001797285, and *Coxiella*-LEs CLEOmar (Fig 5) and AB428 (Fig S16). When the PAI region from *C. burnetii* is compared to *Coxiella*-LEs CLEOmar and AB428, it is completely reshuffled (Fig S16). Contig edges of both CLEOmar and AB428 correspond, most of the time, to IS. However, its order is unclear due to the draft status of their genomes. Furthermore, the PAI region can be detected in *Coxiella*-LEs CRt and CRS-CAT but has been almost totally eroded in CLEDm and *Coxiella*-LEs from *Amblyomma* ticks. Yet, the tRNA-Ile (the putative insertion-site of the PAI) and a few other genes remain (e.g. *rpoD, dnaG*) in CLEDm and *Coxiella*-LEs from *Amblyomma* ticks (Fig 5). Therefore, it is possible that erosion and the inactivation of the PAI are mediated by IS mobilization. This suggests that the PAI was ancestral to the divergence of *C. burnetii* and *Coxiella*-LEs clades, but also that it is no longer required for the mutualistic relationship established by *Coxiella*-LEs within ticks.

### Commonalities and Particularities of pH Homeostasis in the ***C. burnetii*** Lineage

Among the Coxiellaceae, only *C. burnetii* has been reported as acidophilic, a critical trait for its pathogenic cycle (Schaik et al., 2013). Knowing which mechanisms are common to all Coxiellaceae and which are specific to *C. burnetii* could help to understand its pathogenic lifestyle. One proposed specific adaption of acidophilic bacteria concerns the modification of their proteomes, through the enrichment of proteins in basic residues. Hence, it is expected that proteomes from acidophilic bacteria present lower average isoelectric point (pI) than non-acidophilic ones (Baker-Austin and Dopson, 2007). We found significant differences in the average pI among *Coxiellaceae* (Kruskal-Wallis test, *χ*^2^ = 1916.1, *df* = 24, *p − value <* 2.2*e −* 16). Yet, the average pI of proteomes from *C. burnetii* (8.2 *±* 1.9 SD) was similar to other *Coxiellaceae*, including *R. grylli* (8.3 *±* 1.7 SD) or *R. isopodorum* (8.1 *±* 1.7 SD) (Pairwise Wilcoxon Rank Sum tests, *p − value >* 0.05 with false discovery rate adjustment), which are considered non-acidophilic symbionts (Fig S17, Table S10, Supplementary Data). Another acid stress adaptation is the modification of the cell membrane composition, for example, by increasing the percentage of long-chained mono-unsaturated fatty acids (dehydratase FabA) or by synthesizing cyclopropane fatty acids (CFA) (Lund et al., 2014) (Fig 6, Table S11). Only the former mechanism is present in *C. burnetti*, suggesting a possible role of FabA in acid resistance.

**Figure 6.**
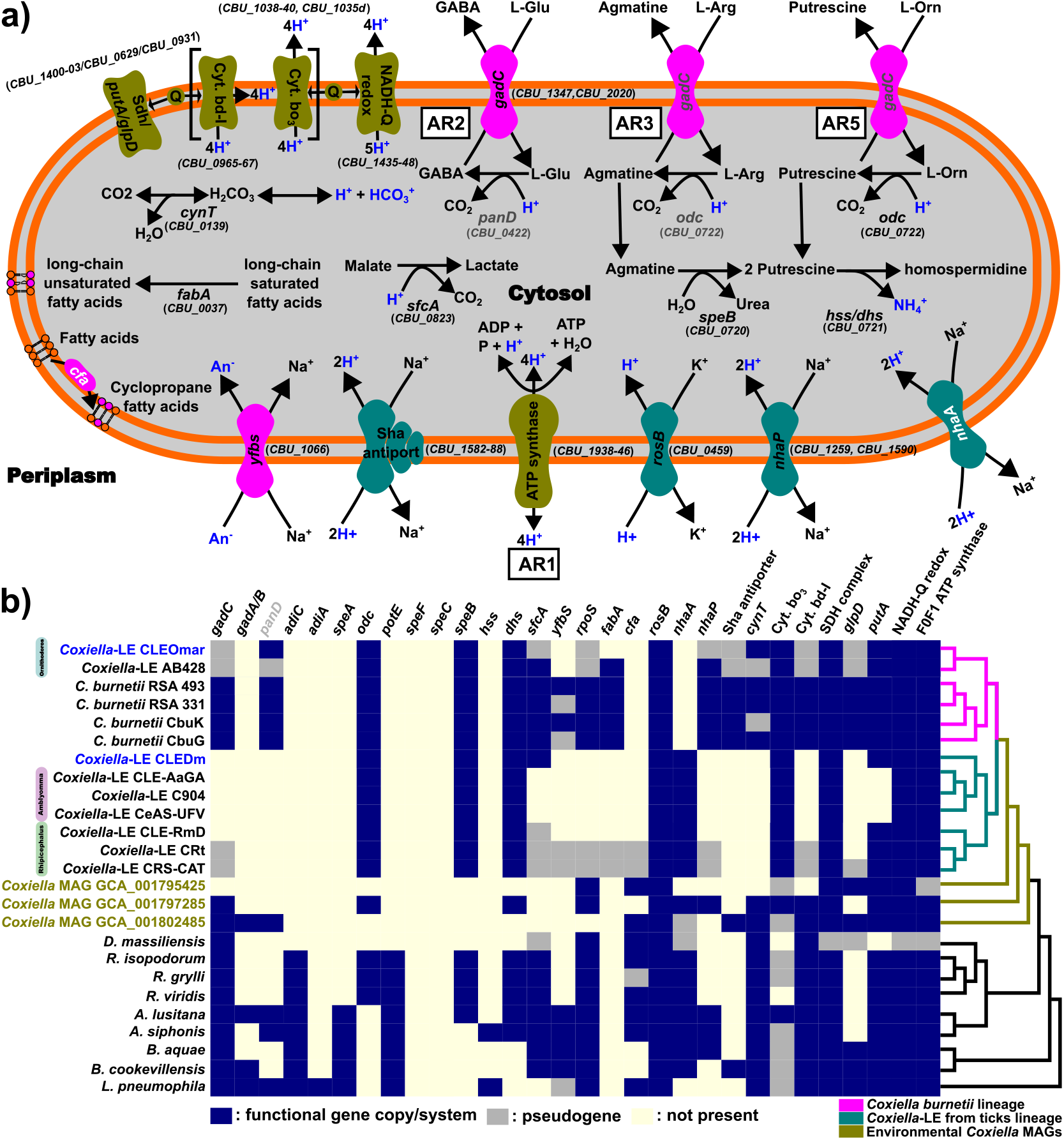
Putative pH regulation mechanisms encoded by *C. burnetii* RSA 493 (**a**)) and their presence in other Coxiellaceae (**b**)). Gene names displayed in gray represents alternative steps probably conducted by co-opted enzymes. Gene names displayed in white represent components not encoded in *C. burnetii* but present, or inactive, in *Coxiella*-LE species. Acid-resistance systems are colored in fuchsia, alkali-resistance in teal blue, and components working in both kind of resistance are displayed in olive green. The cladogram on the right represents the phylogenetic relationships of Coxiellaceae species based on Fig 1. Species color coding is as in Fig 1. Species color coding is as in Fig 1.

In addition, different common mechanisms can help to alleviate acid stress by buffering or extruding H^+^ (Fig 6, Table S11). Among them, acid-resistant (AR) systems play a major role in counteracting acid stress. The AR1 system involves the F1F0-ATPase and other components of the electron transport chain and is present in almost all Coxiellaceae. Likewise, three amino acid-based AR systems were detected among Coxiellaceae: AR2 (glutamate), AR3 (arginine), and AR5 (ornithine). Amino acid-based AR systems import an amino acid molecule, by a specific amino acid antiporter, which is used by a decarboxylase as an H^+^ receptor (Fig 6). AR2 is considered the most efficient AR system and comprises the glutamate antiporter GadC and the glutamate decarboxylase GadB. Most Coxiellaceae encode at least one *gadC* copy, but no *gadB*. Indeed, *C. burnetii* encodes two *gadC*-like transporters (CBU_1347 and CBU_2020), but no *gadB* homolog (Fig 6, Table S11). Since AR2 has been confirmed experimentally as the most important AR in *C. burnetii* (Hackstadt and Williams, 1983), some other decarbolixases may have replaced the GadB function. Among all decarbolixases encoded by *C. burnetii*, only the aspartate 1-decarboxylase PanD seems to be a candidate for replacing the function of GadB: its substrate is close enough to glutamate, and it is present in the *C. burnetii* lineage but absent in almost all Coxiellaceae (Fig 6 b). Finally, no traces of *gadC* or *gadB* genes were detected in most *Coxiella*-LEs (Fig 6 b), except a pseudogenized copy of *gadC* in CLEOmar, AB428, CRt, and CRS-CAT. This pattern suggests that the AR2 system is not required by *Coxiella*-LEs.

Bacteria often need to face environments where the pH is higher than their cytoplasm. In acidophilic bacteria, small increases in external pH can distort their membrane potential, thus, requiring tight control of both cations and anions (Baker-Austin and Dopson, 2007). In such a context, cation antiporters play an important role. Indeed, *C. burnetii* encodes several cation antiporters which are shared with other Coxiellaceae (Fig 6, Table S11). Already mentioned above, the Na^+^/H^+^ Sha/Mrp antiporter, was acquired laterally by the MRCA of the *C. burnetii* lineage (Fig S15-S15). Besides their role as cation antiporters, Sha/Mrp antiporters have other functions than pH homeostasis (Ito et al., 2017), including virulence and host-colonization (Kosono, Haga, et al., 2005).

## Discussion

### The Parasitism-Mutualism Continuum in ***Coxiella***

The Coxiellaceae family is mainly composed of bacteria found in aquatic environments or associated with arthropods or amoebae and involved in different symbiotic relationships. Some species, such as *Aquicella* and *Berkiella*, are facultative parasites of aquatic amoebae (Mehari et al., 2016; Santos et al., 2003). Others, such as *R. viridis*, are defensive symbionts in aphids (Łukasik et al., 2013). Based on their genome size, *Coxiella* MAGs could range from free-living aquatic bacteria (genomes generally larger than 4 Mb) to endosymbionts (genomes usually below 1.5 Mb) (Latorre and Manzano-Marín, 2017; Moran and Bennett, 2014). Within the *Coxiella* genus, two clear examples of opposite symbiotic relationships are *C. burnetii* and *Coxiella*-LEs. While the first is an obligate parasite of mammals (Voth and Heinzen, 2007), the latter are considered obligatory mutualistic symbionts of ticks (Duron and Gottlieb, 2020). Based on the monophyly of *C. burnetii* and all *Coxiella*-LEs, it was suggested that the former arose from a mutualistic tick symbiont that acquired virulence (Borucinska, 2016; Duron, Noël, et al., 2015).

To examine this question further, Brenner et al. (2021) sequenced a *Coxiella*-LE associated with the soft tick *Ornithodoros amblus*, a close relative of *C. burnetii*, although more derived than CLEOmar. Indeed, CLEOmar is the basal species of the clade containing *Coxiella*-LE from *O. amblus* (Brenner et al., 2021). In their work, Brenner et al. (2021) proposed that mutualistic *Coxiella*-LEs found in ticks derived from a parasitic ancestor able to invade different hosts. Their conclusion was based on several phylogenetic and comparative genomic analyses: **(i)** the monophyletic origin of *C. burnetii* and *Coxiella*-LEs after increasing the taxon sampling compared to previous works (Duron, Noël, et al., 2015); **(ii)** the presence of several pathogenic bacteria across *Coxiella*-LE lineages; **(iii)** the universal presence of the Dot/Icm T4SS across Coxiellaceae, except in *Coxiella*-LEs for which it is pseudogenized; **(iv)** the fact that *Coxiella*-LEs are a streamlined version of *C. burnetii*, including cell walls, pH regulation, free-radical protection, and antimicrobial transporters, which are crucial elements for pathogenic lifestyles; **(v)** most gene acquisitions took place in the *C. burnetii*/*Coxiella*-LEs MRCA suggesting an inability of *Coxiella*-LEs to acquire new genetic material (Brenner et al., 2021).

Using an expanded set of Coxiellaceae species including environmental species and two new *Coxiella*-LEs, our results are in agreement with those presented by Brenner et al. (2021). Our phylogenomic tree corroborates the monophyly of *C. burnetii*/*Coxiella*-LEs, but also suggests the clade originated from aquatic bacteria able to establish symbiotic relationships with different hosts. We also observed that *Coxiella*-LEs are a reduced version of *C. burnetii* with few species-specific genes. (Gottlieb et al., 2015) proposed that *Coxiella*-LEs do not face the acidic environment of the *Coxiella* Containing Vacuole (CCV), a key feature of *C. burnetii*, because they are harbored inside host-derived vacuoles (Klyachko et al., 2007; Lalzar, Friedmann, et al., 2014), which are thought to be more pH neutral. As also found by Brenner et al. (2021), we found traces of acid-resistance systems (*sfcA, gadC*, and *cfa*) in CLEDm and some *Coxiella*-LE from *Rhipicephalus* tick species. Their presence suggests that the *C. burnetii*-*Coxiella*-LE MRCA was probably a pathogen able to confront acidic environments.

It has been proposed that *Coxiella*-LEs play a critical role in tick development as they supply different B vitamins and co-factors lacking from the blood-based diet of ticks (Duron and Gottlieb, 2020; Manzano-Marín, Oceguera-Figueroa, et al., 2015). In their work, Brenner et al. (2021) suggested that the production of B vitamins and co-factors triggered the evolution of the *C. burnetii*/*Coxiella*-LEs MRCA towards more mutualistic interactions with ticks. We found that these biosynthetic pathways are also encoded in many environmental *Coxiella* MAGs, including those closely related to the *C. burnetii*/*Coxiella*-LE clade, supporting their presence in the *Coxiella* MRCA. Interestingly, B vitamin and co-factor biosynthesis pathways are also found in pathogenic Coxiellaceae such as *Rickettsiella, Aquicella*, and *Berkiella*. Biotin synthesis is likewise critical for virulence in some human pathogens, such as *Francisella tularensis* and *Mycobacterium tuberculosis*, and may have played similar roles in parasitic Coxiellaceae (Feng et al., 2014; Park et al., 2011). Indeed, blocking biotin biosynthesis inhibits *C. burnetii* growth on specific axenic media (Moses et al., 2017), suggesting that it is required for the normal development of this pathogen. Thus, the potential to synthesize vitamins and co-factors is not a good predictor of mutualistic relationships but rather may have act as a pre-adaptation for establishing interactions with blood-feeding arthropods.

The Dot/Icm T4SS is described as a virulence factor in many pathogenic bacteria and its presence can be considered a signature of pathogenicity (Gomez-Valero, Chiner-Oms, et al., 2019; Schaik et al., 2013). We detected the Dot/Icm T4SS (or signatures of it) in all Coxiellaceae, including the environmental *Coxiella* MAGs, confirming its universal presence across this family (Brenner et al., 2021). While most *Coxiella*-LEs have lost all relevant genes, some pseudogenes are still detectable in *Coxiella*-LEs from *Ornithodoros* soft (Brenner et al. (2021) and this work) and hard ticks lineages (Buysse, Duhayon, et al., 2021; Gottlieb et al., 2015). Similar to Brenner et al. (2021), we conclude that the *C. burnetii*/*Coxiella*-LE MRCA encoded a functional Dot/Icm T4SS and was able to parasitize different hosts, including amoebae (La Scola and Raoult, 2001). In recently established symbionts, the loss of virulence-associated secretion systems, together with increased vertical transmission and restricted tropism can facilitate the switch towards more mutualistic relationships (Manzano-Marín, Simon, et al., 2016; Oakeson et al., 2014; Yamamura, 1993). Therefore, the inactivation of the Dot/Icm T4SS could have facilitated the domestication of facultative parasitic *Coxiella* by ticks.

We identified a genomic region in *C. burnetii* which resembles a pathogenic island (PAI). Generally, PAIs are defined as genomic islands (10-200 Kb) enriched in genes related to virulence, antibiotic resistance, symbiosis, and environmental fitness. PAIs are typically horizontally acquired (exogenous DNA) and often include tRNAs, which act as integration sites for the PAI. The presence of repeats and mobile elements make PAIs dynamic, favoring recombination and gene exchanges, but also leads to rapid gene losses when they are no longer required (Hacker and Kaper, 2000). The putative PAI region in *C. burnetii* contains a tRNA that could have served as an integration point (isoleucine 2 anticodon), several mobile elements (ISs), other repeats, different horizontally acquired genes, the Dot/Icm T4SS (considered a virulence factor) and different effectors associated with it. In *Coxiella*-LEs CLEOmar and AB428, this region is distributed among several contigs flanked by ISs and shows partial synteny with *C. burnetii*. Syntenic regions to the PAI, including the isoleucine tRNA, are also detected in *Coxiella* sp. GCA_001802485 and GCA_001797285 MAGs. However, all four of these genomes are still drafts and the full PAI structure is unknown. Interestingly, a similar PAI-like region, with the same isoleucine tRNA and adjacent genes and with remnants of the Dot/Icm T4SS, was detected in *Coxiella*-LEs CRt and CRSCAT. In the smallest *Coxiella*-LEs, CLEDm and those from *Amblyomma* ticks, only the putative insertion site of the PAI, the isoleucine tRNA and adjacent genes, are maintained. Assuming the most parsimonious scenario, our results support a single acquisition of the PAI by the *C. burnetii*/*Coxiella*-LEs MRCA and its progressive loss in non-pathogenic *Coxiella*-LEs. *Coxiella*-LEs CLEOmar and AB428 are, like other recent host-associated symbionts, overrun by mobile elements (Latorre and Manzano-Marín, 2017). Their PAI-like and adjacent regions could therefore have been reshuffled and many genes inactivated by the activity of mobile elements. This could have resulted in the rapid loss of functions no longer required by the symbiont, such as the Dot/Icm T4SS or the Sha (Hacker and Kaper, 2000; Latorre and Manzano-Marín, 2017). Thus, we can speculate that the fast inactivation of genes related to virulence, pathogenicity, or environmental response might have played a significant role during the transition towards more mutualistic interactions in *Coxiella*-LEs (Yamamura, 1993).

In summary, the *C. burnetii*/*Coxiella*-LEs MRCA was likely more similar to *C. burnetii*: an acidophilic pathogen with a putative PAI encoding a virulence factor (the Dot/Icm T4SS) and other functions related to environmental fitness (H^+^ antiporters). The ability to produce B vitamins and co-factors, together with a reduction in its virulence, could have aided *Coxiella* bacteria to evolve towards more mutualistic interactions in some lineages.

### *C. burnetii* Specialization: Acid Resistance and Biphasic Life Cycle

*C. burnetii* is the only known acidophilic bacteria (Hackstadt, 1983) among Legionellales pathogens. It presents three amino acid Acid Resistance (AR) systems that work in a pH range of 4-6, close to that of the different *Coxiella* Containing Vacuole (CCV) phases (Foster, 2004). The glutamate system (AR2) is the most effective (Lund et al., 2014) with optimal AR2 decarboxylase activity at pH 4, close to the CCV pH (Foster, 2004). Interestingly, *C. burnetii* encodes two GadC-like transporters, *CBU_1347* and *CBU_2020*. According to the TCDB classification engine, *CBU_1347* presents the highest identity to GadC from *Escherichia coli* (GenBank: BAI30440.1). In addition, *CBU_1347* is up-regulated during the transition from SCV to LCV (Sandoz et al., 2016), suggesting it may be the main AR2 antiporter. It may be that additional GadC copies in *C. burnetii* and *Coxiella* MAGs provide a broader substrate range by assuming the function of AdiC (AR3) or PotE (AR5).

Almost no Coxiellaceae, including *C. burnetii*, encode the AR2 decarboxylase cognate GadB, which uses the negatively charged amino acid L-glutamate as proton receptor. However, *C. burnetii*, CLEOmar, several *Coxiella* MAGs, and *Aquicella* species encode an aspartate 1-decarboxylase PanD which may decarboxylate L-aspartate, another negatively charged amino acid (Williamson and Brown, 1979). This enzyme presents the lowest pI (4.7) in *C. burnetii* (CBU_0422) when compared to almost all other PanD from the Coxiellaceae. Indeed, its pI and predicted charge at 5.5 pH (the pH of the lysosome) are closer to those of GadB from some acidophile bacteria (Table S12). Therefore, we propose that PanD might have been co-opted to work as part of the AR2 system by decarboxylating L-glutamate under acidic conditions (Kelkar and Ochman, 2013). However, the ability of PanD from *C. burnetii* to perform in acidic environments should be empirically validated.

Similarly, when Coxiellaceae genomes encoding the AR3 are compared, the ornithine decarboxylase Odc and the arginine decarboxylase SpeA present a mutually exclusive pattern, suggesting that ornithine decarboxylase may provide a broader substrate range and decarboxylate both arginine (AR3) and ornithine (AR5). Co-opting enzymes, such as PanD or Odc, can result as a consequence of the genome reduction process and the trend to minimize functional redundancy in symbionts (Manzano-Marín, Oceguera-Figueroa, et al., 2015; Murray et al., 2020). That being the case, AR systems from *C. burnetii* and other Coxiellaceae species could be based on the co-option.

Interestingly, *C. burnetii* uses glutamate as a primary energy source within an effective range between 2 to 5.5 pH, a range similar to the phagosomes/CCV (Hackstadt and Williams, 1981, 1983; Omsland et al., 2008). The phagosome may present low nutrient levels and using non-essential amino acids as an energy source is a common adaptation in pathogens (Omsland et al., 2008). Therefore, it might be that the glutamine present in the SCV phase-specific ScvA protein (*∼* 23%) can be converted directly to glutamate for energetic purposes (catabolism) or pH regulation (decarboxylation) during the early phagosome invasion. If so, *C. burnetii* can overcome acid stress without the need to scavenge glutamate from its host. In this context, we can think of ScvA as a unique adaptation of *C. burnetii* to pathogenicity, where ScvA plays a role in both SCV formation (Minnick and Raghavan, 2012) and acid-resistance.

Another particularity of *C. burnetii* is that it presents a biphasic life cycle (Coleman, Fischer, Howe, et al., 2004; Coleman, Fischer, Cockrell, et al., 2007; Minnick and Raghavan, 2012; Schaik et al., 2013; Voth and Heinzen, 2007). The PAI of *C. burnetii* presents a Sha/Mrp antiporter (organized as an operon) located *∼*25 Kb upstream from the Dot/Icm T4SS. The Sha/Mrp is a Na^+^/H^+^ antiporter involved in the establishment and maintenance of Na^+^ electrochemical potential, extrusion of Na^+^/Li^+^ for avoiding toxic concentrations, cell volume regulation, and pH maintenance under alkaline stress. Also, it has been shown to play several roles in addition to pH homeostasis (Ito et al., 2017). While in *Bacillus subtilis* the disruption of *shaA* resulted in sporulation-deficient phenotypes (Kosono, Ohashi, et al., 2000), in *P. aeruginosa* strain PAO1 it reduced bacterial virulence and colonization capabilities (Kosono, Haga, et al., 2005). The Sha/Mrp antiporter seems to also play an important function in establishing the *Rhizobium meliloti*-plant symbiotic relationship, where only symbionts able to grow in alkaline environments can colonize the plant roots (Putnoky et al., 1998). Our results suggest that the *sha* operon was acquired by the MRCA of the *C. burnetii* lineage. However, it is not clear if the acquisition was from a close relative, such as *Coxiella* sp. GCA_001802485 MAG, or more directly from a *Legionella* bacterium. Based on the reported functions of the Sha/Mrp antiporter, we propose that in *C. burnetii* it is not only related to alkali resistance but could also be involved in the SCV formation, hence, pathogenesis. If this is so, the inactivation of the Sha/Mrp antiporter may produce attenuated phenotypes in *C. burnetii* as in *P. aeruginosa*, but may also limit transmission and dispersal potential by compromising SCV formation (Kosono, Haga, et al., 2005).

### Concluding Remarks

Based on comparative genomic approaches using Coxiellaceae species with different lifestyles, ranging from free-living to obligate mutualist symbionts, we propose a scenario for the origin of mutualistic *Coxiella* endosymbionts in ticks. An environmental, and probably pathogenic, *Coxiella* ancestor invaded different hosts, thanks to the presence of the Dot/Icm T4 Secretion System, and other genes, included in a putative pathogenic island and adjacent regions. This ancestor evolved into two lineages, one including *C. burnetii* and the other including mainly tick-associated symbionts. The ability of the *Coxiella*-LE ancestor to produce B vitamins and co-factors contributed to its domestication in some tick species, evolving later on towards more mutualistic symbiosis.

A more recent process of transition towards mutualism from parasitism can be observed within the *C. burnetii* lineage. In this lineage, its ancestor laterally acquired a Sha/Mrp antiporter close to the Dot/Icm region. Based on previously reported functions of the Sha/Mrp antiporter, we hypothesize that its acquisition might have enabled *C. burnetii* to resist alkaline environments found outside the host. Moreover, the Sha operon might be involved in the development of the Small-Cell Variant resistant form of *C. burnetii* (Ito et al., 2017; Kosono, Ohashi, et al., 2000). In *Coxiella*-LE CLEOmar and AB428, members of the *C. burnetii* lineage, the Dot/Icm T4SS, the Sha/Mrp antiporter, and the acid-resistance systems are inactive, or almost inactive. Their combined inactivation probably reduced the virulence, dispersion, and tropism of the CLEOmar ancestor thereby increasing the benefits for the host harboring a symbiont able to supplement its diet with B vitamins and co-factors (Manzano-Marín, Oceguera-Figueroa, et al., 2015; Manzano-Marín, Simon, et al., 2016). Selection would then have increased the vertical transmission of the symbiont, aligning both host and pathogen fitness, thus facilitating the emergence of mutualism in CLEOmar (Yamamura, 1993). As the Dot/Icm T4SS is widespread in the *Coxiella* genus, it could have allowed them to exploit different hosts, such that, the emergence of mutualistic representatives could occur on multiple occasions in the *Coxiella* genus, as is the case for CLEOmar and *Coxiella*-LEs which belong to different lineages.

## Acknowledgments

DSG was supported by the European Union’s Horizon 2020 research and innovation program under a Marie Skłodowska-Curie Individual Fellowship (GuardSym, grant agreement no. 885583). We would like to acknowledge the whole PCI Community for their effort and time, especially Daniel Tamarit, Sophie Abby, Adam Ossowicki, and one anonymous reviewer for their constructive suggestions.

## Data, Script, Code, and Supplementary Information Availability

RAW reads generated for this work and *Coxiella*-LE CLEOmar (GCA_907164965) and CLEDm (GCA_907164955) genome assemblies are available European Nucleotide Archive (ENA) under the BioProject number PRJEB44453. *Coxiella*-LE CLEOmar and CLEDm annotated genomes, relevant performed analysis, scripts, and used data are available FigShare https://doi.org/10.6084/m9.figshare.12563558.v3. All phylogenetic trees can be accessed at https://itol.embl.de/shared/dsantosgarcia

