## Supplementary Figures for "Genomic Changes During the Evolution of the *Coxiella* Genus Along the Parasitism-Mutualism Continuum"

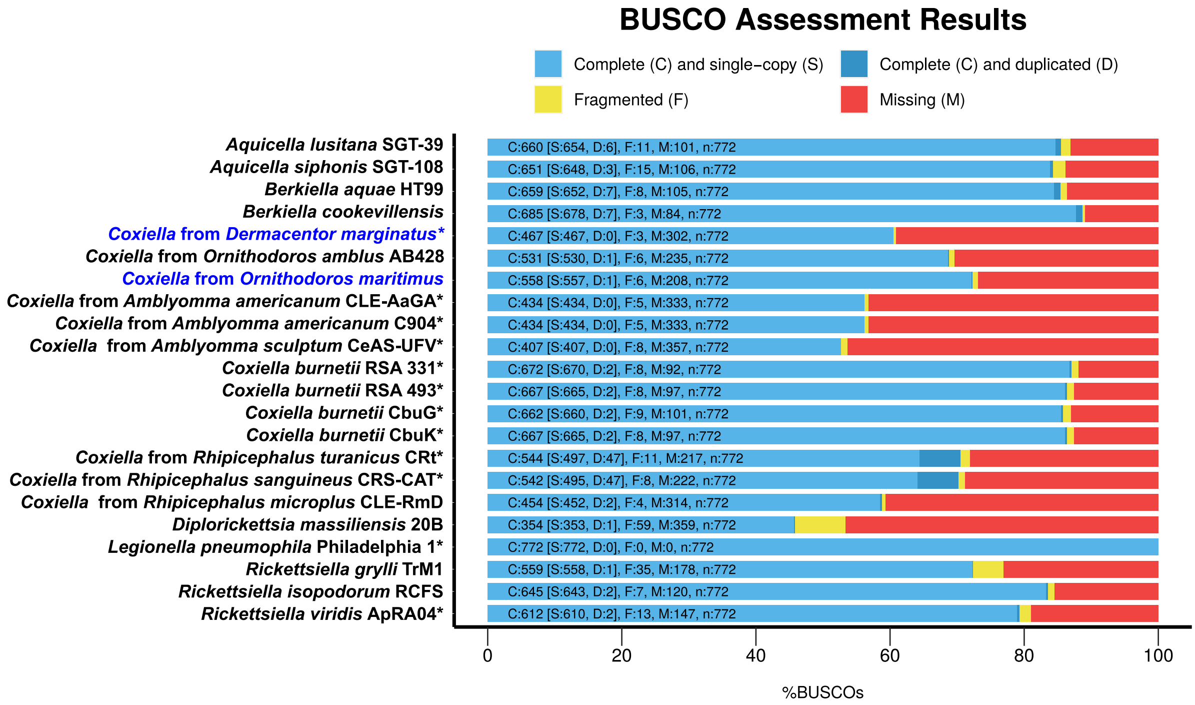


**Figure S1.** Completeness of Coxiellacea genomes assessed using BUSCO pipeline. *Coxiella*-LE genomes obtained in this work are highlighted in blue. *Legionella pneumophila* was used as an outgroup. *: closed circular genomes.

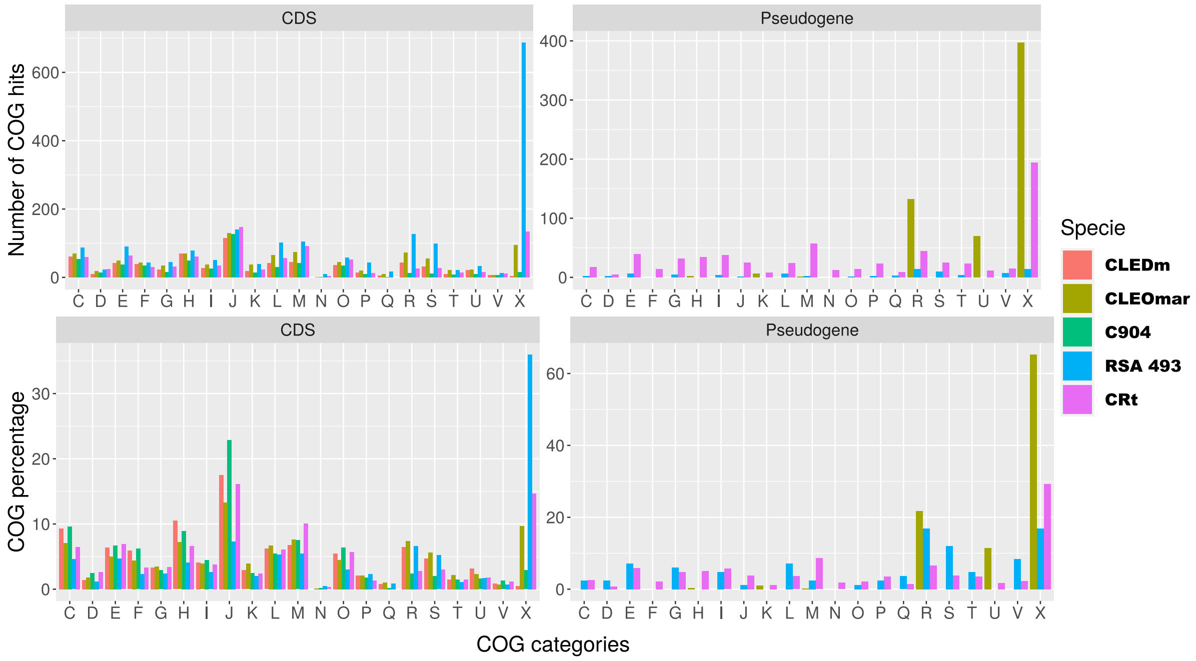


**CELLULAR PROCESSES AND SIGNALING**

**[D] Cell cycle control, cell division, chromosome partitioning**
**[M] Cell wall/membrane/envelope biogenesis**
**[N] Cell motility**
**[O] Post-translational modification, protein turnover, and chaperones**
**[T] Signal transduction mechanisms**
**[U] Intracellular trafficking, secretion, and vesicular transport**
**[V] Defense mechanisms**
**[W] Extracellular structures**
**[Y] Nuclear structure**
**[Z] Cytoskeleton**

**INFORMATION STORAGE AND PROCESSING**

**[A] RNA processing and modification**
**[B] Chromatin structure and dynamics**
**[J] Translation, ribosomal structure and biogenesis**
**[K] Transcription**
**[L] Replication, recombination and repair**

**METABOLISM**

**[C] Energy production and conversion**
**[E] Amino acid transport and metabolism**
**[F] Nucleotide transport and metabolism**
**[G] Carbohydrate transport and metabolism**
**[H] Coenzyme transport and metabolism**
**[I] Lipid transport and metabolism**
**[P] Inorganic ion transport and metabolism**
**[Q] Secondary metabolites biosynthesis, transport, and catabolism**

**POORLY CHARACTERIZED**

**[R] General function prediction only**
**[S] Function unknown**
**[X] No hits**

**Figure S2.** Comparison between COG assignment (top bar-plots) and their respective percentage (bottom) for coding sequences (CDS) and pseudogenes found in *Coxiella burnetii* RSA 493 (RSA 493) and *Coxiella*-LEs from *Rhipicephalus turanicus* (CRt), *Dermacentor marginatus* (CLEDm), *Ornithodoros maritimus* (CLEOmar), and *Amblyomma americanum* C904 (C904). COG categories with less than five hits were not plotted.

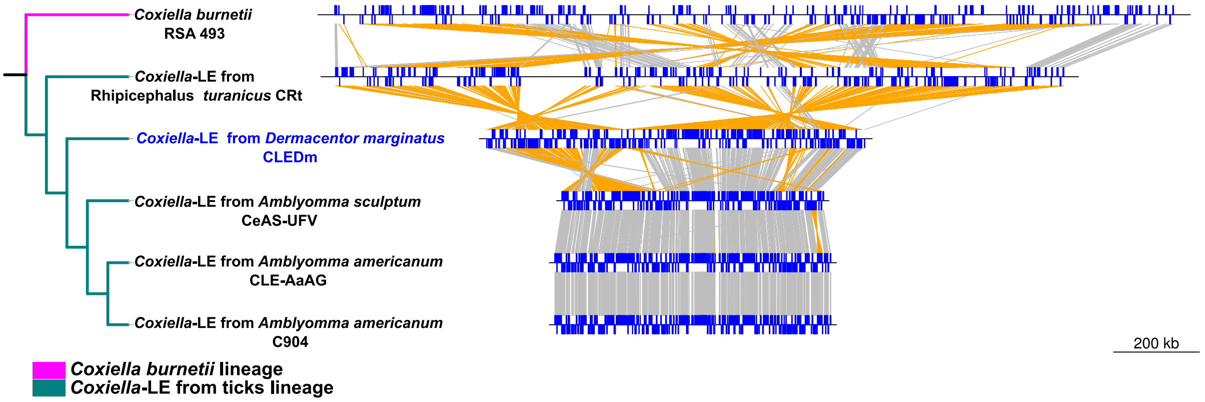


**Figure S3.** Genomic synteny based on 458 single copy shared genes between *Coxiella brunetii* RSA 493 and *Coxiella*-LEs from *Rhipicephalus*, *Dermacentor* (highlighted in blue, obtained for this work), and *Amblyomma* tick species. Genomes are represented as black lines, shared genes in the direct and complementary strand are displayed as upwards and downwards blue boxes. Gray lines connect genes in the same strand, while yellow lines connect genes in different strands. Twisted lines indicate inversions. The cladogram on the left represents the phylogenetic relationships of Coxiella species based on Fig 1. Species color coding is as in Fig 1.

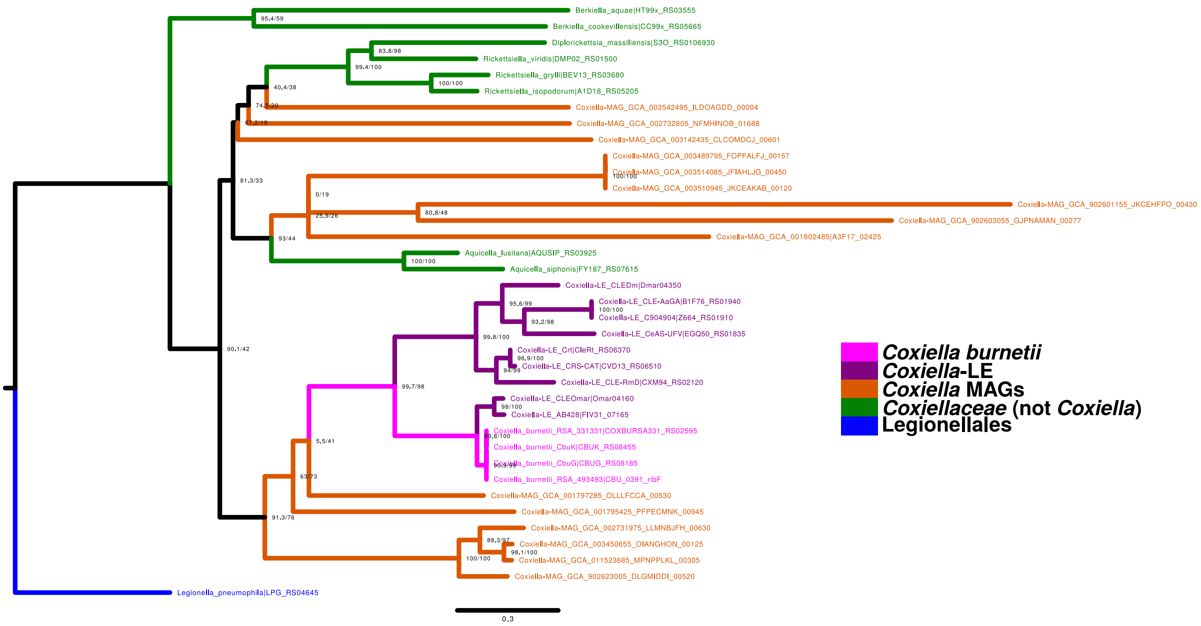


**Figure S4.** Riboflavin kinase RifF Clusters of Orthologous Proteins (COPS) Maximum Likelihood phylogenetic tree. The numbers at each node represent 1,000 SH-aLRT (left) and ultrafast bootstrap (right) support. *L. pneumophila* was set as outgroup.

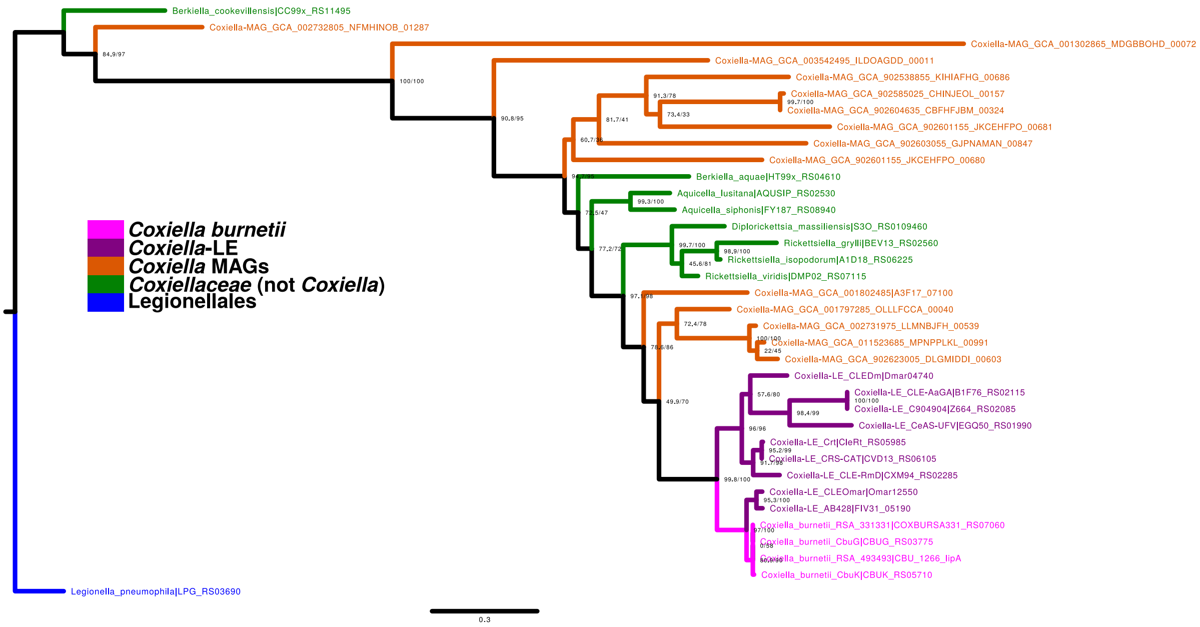


**Figure S5.** Lipoyl synthase LipA Clusters of Orthologous Proteins (COPS) Maximum Likelihood phylogenetic tree. The numbers at each node represent 1,000 SH-aLRT (left) and ultrafast bootstrap (right) support. *L. pneumophila* was set as outgroup.

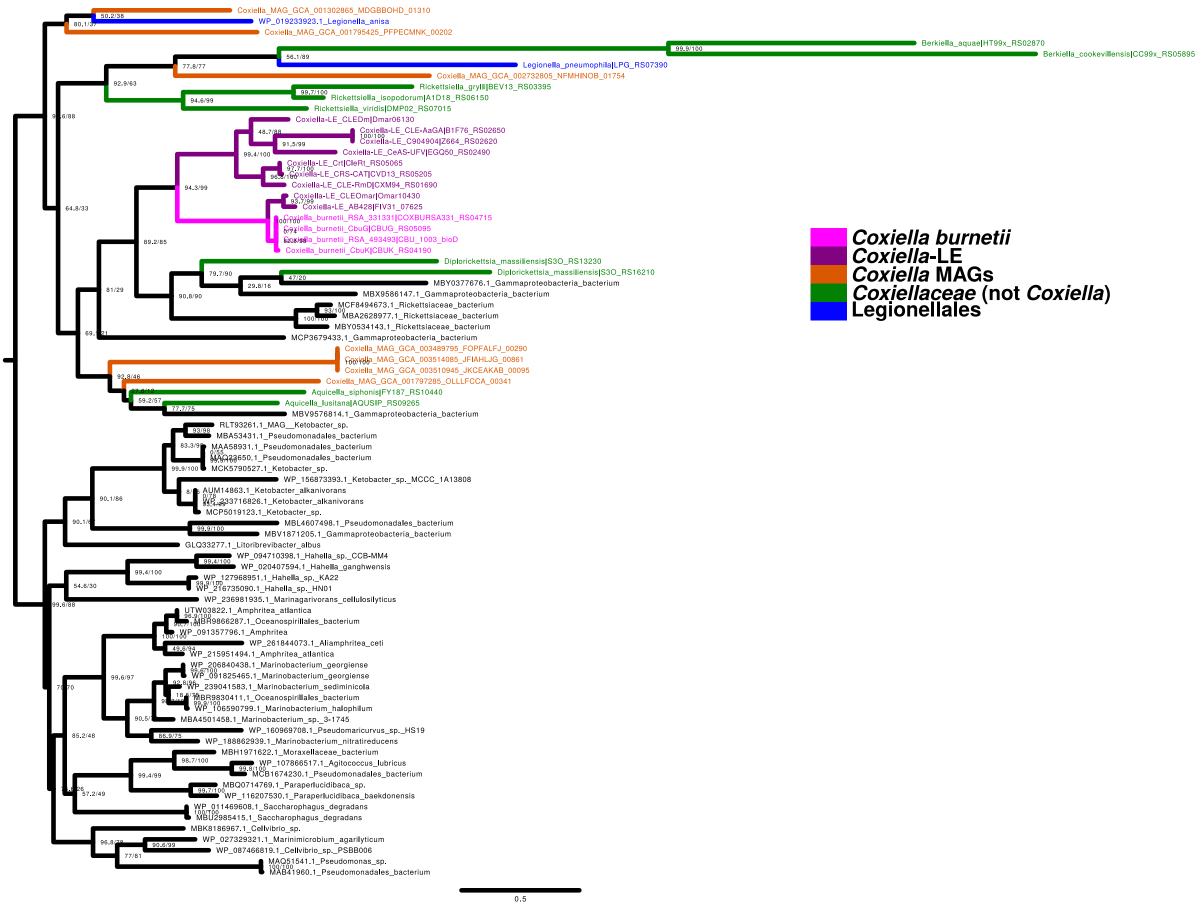


**Figure S6.** Mid-point rooted Maximum Likelihood phylogenetic tree of dethiobiotin synthetase BioD. Obtained BioD Cluster of Orthologous Proteins (COPS) together with the first 50 BLASTP hits (non-redundant database, excluding *Coxiella* species) were included in the reconstruction. The numbers at each node represent 1,000 SH-aLRT (left) and ultrafast bootstrap (right) support.

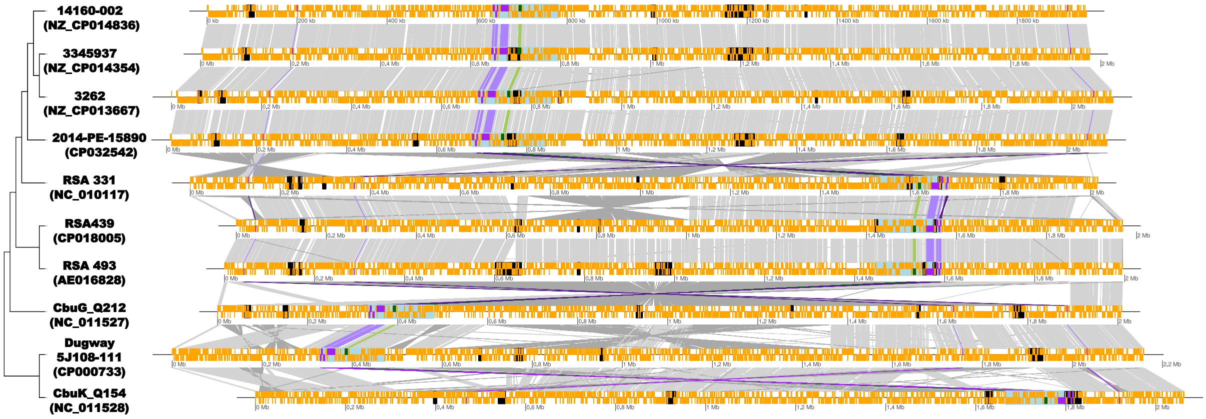


**Figure S7.** Whole genome synteny among different *C. burnetii* strains. The 144 kb region containing the PAI, which includes the Dot/Icm T4SS, and the Sha/Mrp antiporter is highlighted in pale blue. A 6.5 Kb inversion containing four Dot/Icm T4SS genes (*dotA*, *dotB*, *icmX*, and *icmV*) is present in RSA 331 strain. Gray lines connect orthologous genes. Inversions are denoted by dark gray twisted lines. Dot/Icm T4SS genes are highlighted in purple. Dark purple lines indicate Dot/Icm T4SS genes that has suffered an inversion. Sha/Mrp antiporter genes are highlighted in green. Black boxes denote identified genomic islads by IslandView 4 (Table S8). The cladogram in the left represents *C. burnetii* strains phylogenetic relationships. Accession numbers are displayed between parentheses.

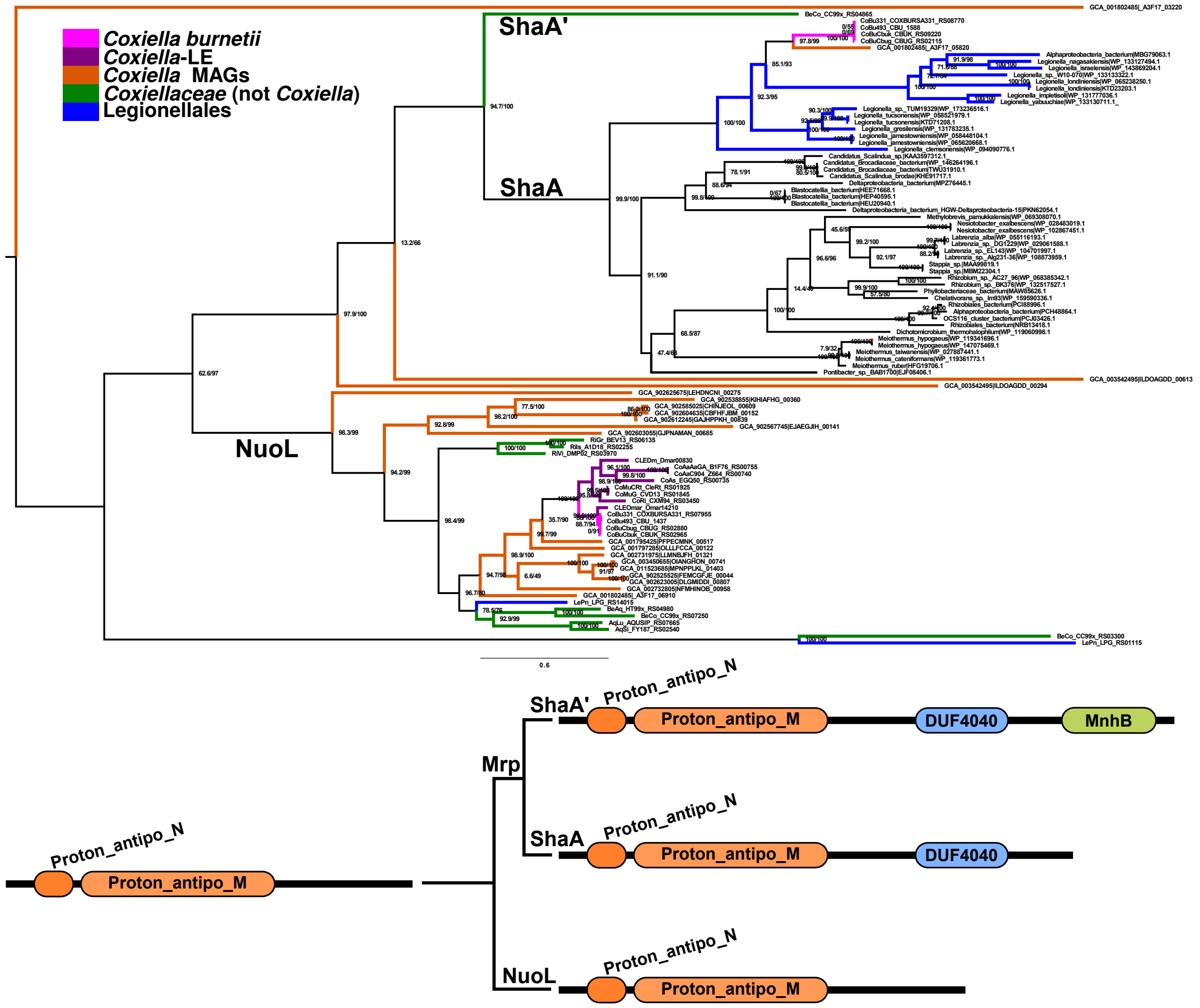


**Figure S8.** Mid-point rooted Maximum Likelihood phylogenetic tree of ShaA and NuoL (top). Obtained ShaA and NuoL Clusters of Orthologous Proteins (COPS) together with the first 50 BLASTP hits (non-redundant database, excluding *Coxiella* species) were included in the reconstruction. The green box denotes ShaA encoded by *Coxiella burnetii* strains and the basal *Coxiella* MAG GCA_ 001802485. The numbers at each node represent 1,000 SH-aLRT (left) and ultrafast bootstrap (right) support. Summarized cladogram (bottom) representing the different domains in NuoL, ShaA, and fused ShaA-ShaB (ShaA'). Environmental *Coxiella* MAGs are denoted by their GenBank accession number (starting with GCA).

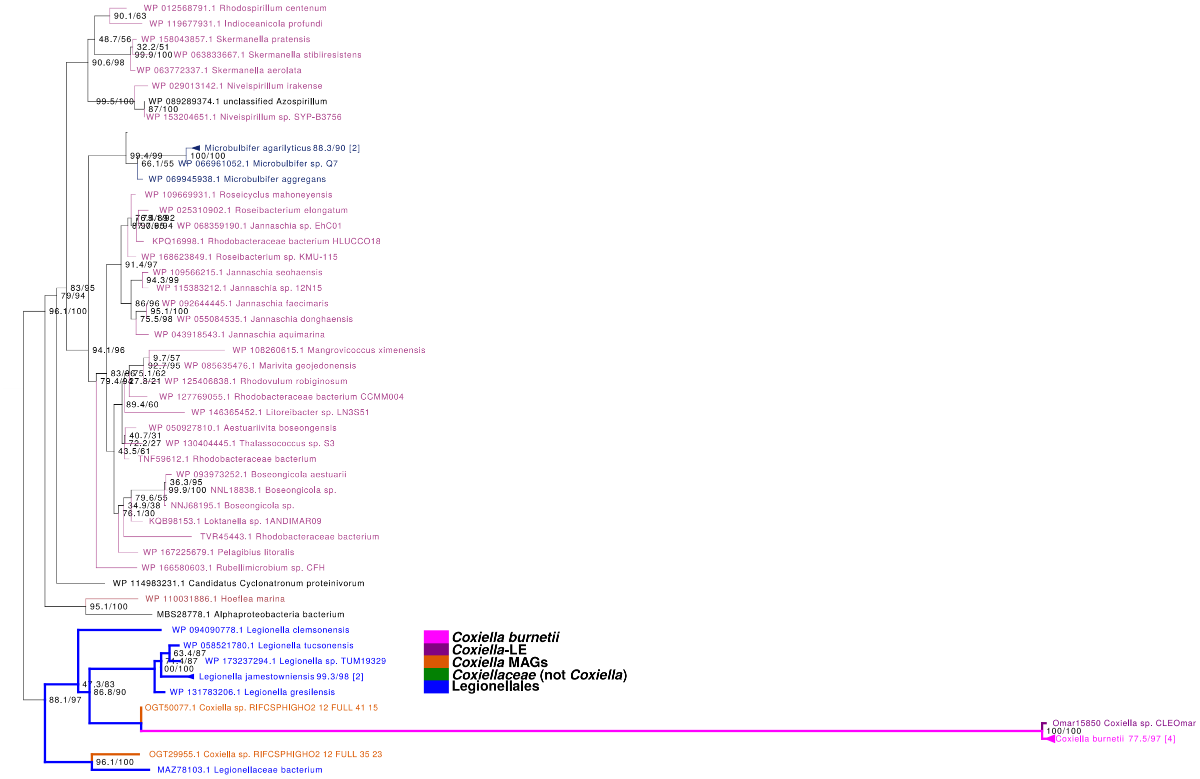


**Figure S9.** Mid-point rooted Maximum Likelihood phylogenetic tree of ShaB. Obtained ShaB Cluster of Orthologous Proteins (COPS) together with the first 50 BLASTP hits (non-redundant database, excluding *Coxiella* species) were included in the reconstruction. The clade containing ShaB from *Coxiella burnetii* is denoted by thicker lines. The numbers at each node represent 1,000 SH-aLRT (left) and ultrafast bootstrap (right) support.

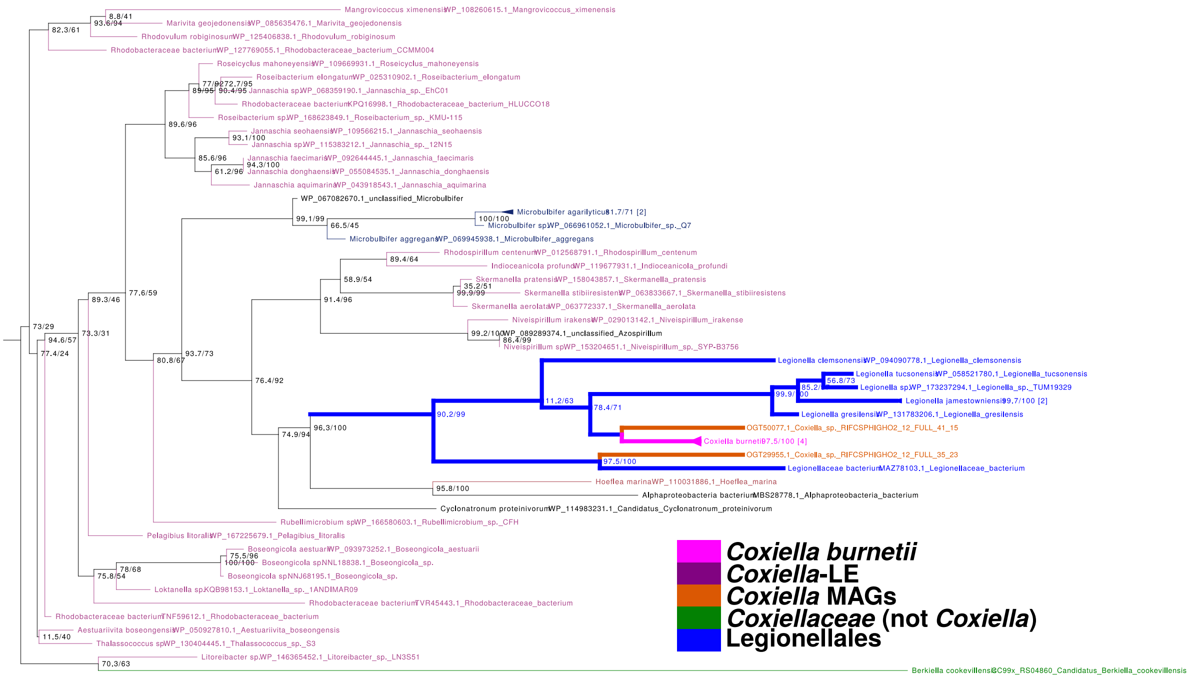


**Figure S10.** Mid-point rooted Maximum Likelihood phylogenetic tree of ShaC. Obtained ShaC Cluster of Orthologous Proteins (COPS) together with the first 50 BLASTP hits (non-redundant database, excluding *Coxiella* species) were included in the reconstruction. The clade containing ShaC from *Coxiella burnetii* is denoted by thicker lines. The numbers at each node represent 1,000 SH-aLRT (left) and ultrafast bootstrap (right) support.

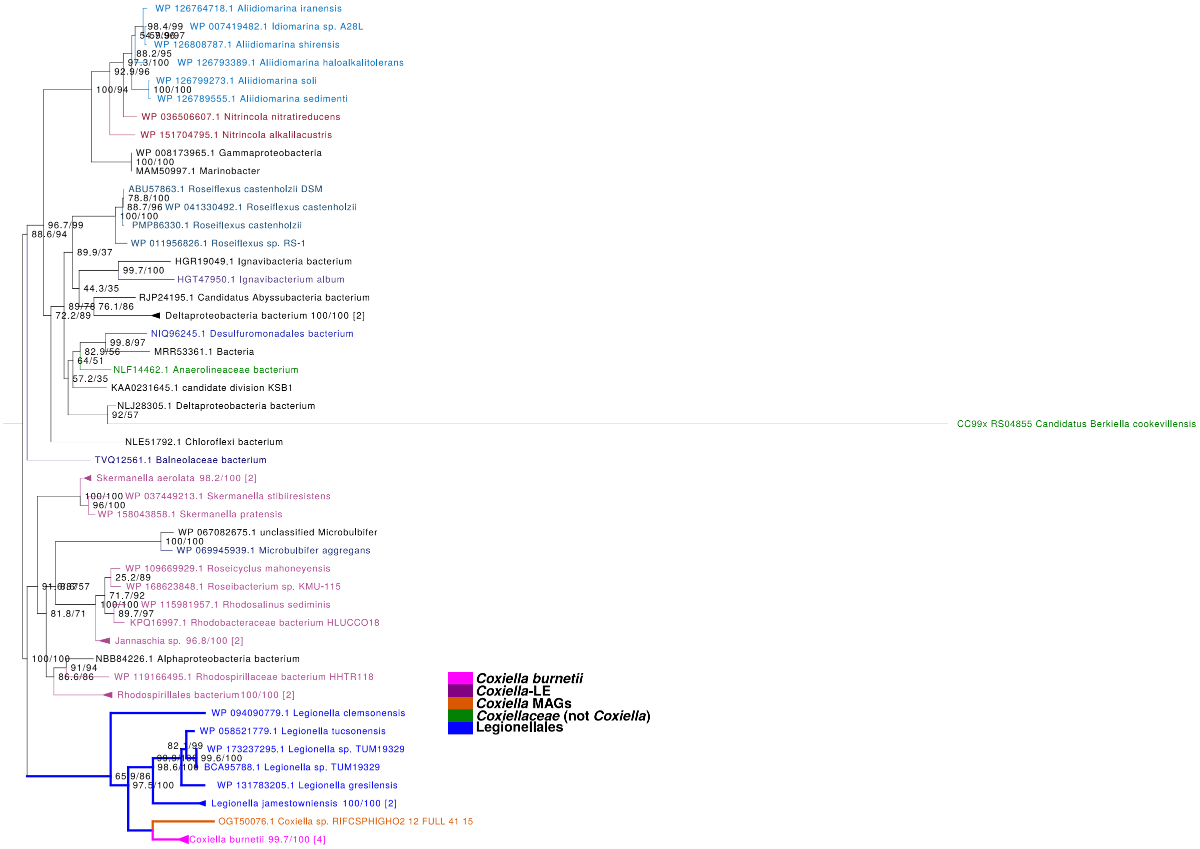


**Figure S11.** Mid-point rooted Maximum Likelihood phylogenetic tree of ShaD. Obtained ShaD Cluster of Orthologous Proteins (COPS) together with the first 50 BLASTP hits (non-redundant database, excluding *Coxiella* species) were included in the reconstruction. The clade containing ShaD from *Coxiella burnetii* is denoted by thicker lines. The numbers at each node represent 1,000 SH-aLRT (left) and ultrafast bootstrap (right) support.

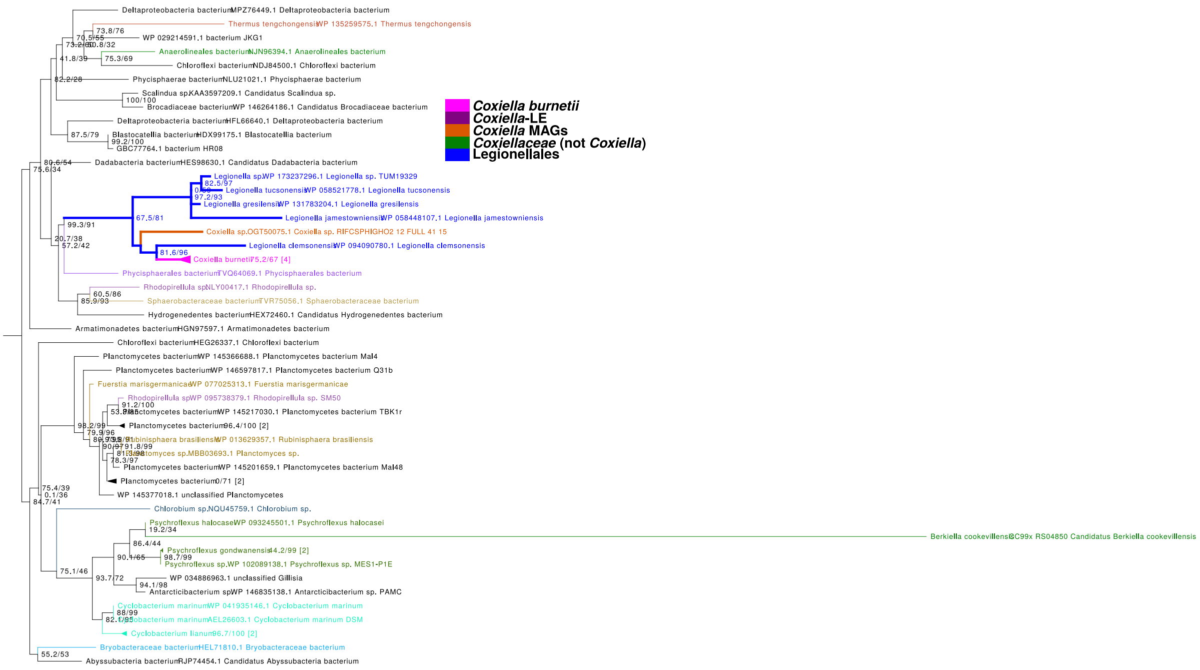


**Figure S12.** Mid-point rooted Maximum Likelihood phylogenetic tree of ShaE. Obtained ShaE Cluster of Orthologous Proteins (COPS) together with the first 50 BLASTP hits (non-redundant database, excluding *Coxiella* species) were included in the reconstruction. The clade containing ShaE from *Coxiella burnetii* is denoted by thicker lines. The numbers at each node represent 1,000 SH-aLRT (left) and ultrafast bootstrap (right) support.

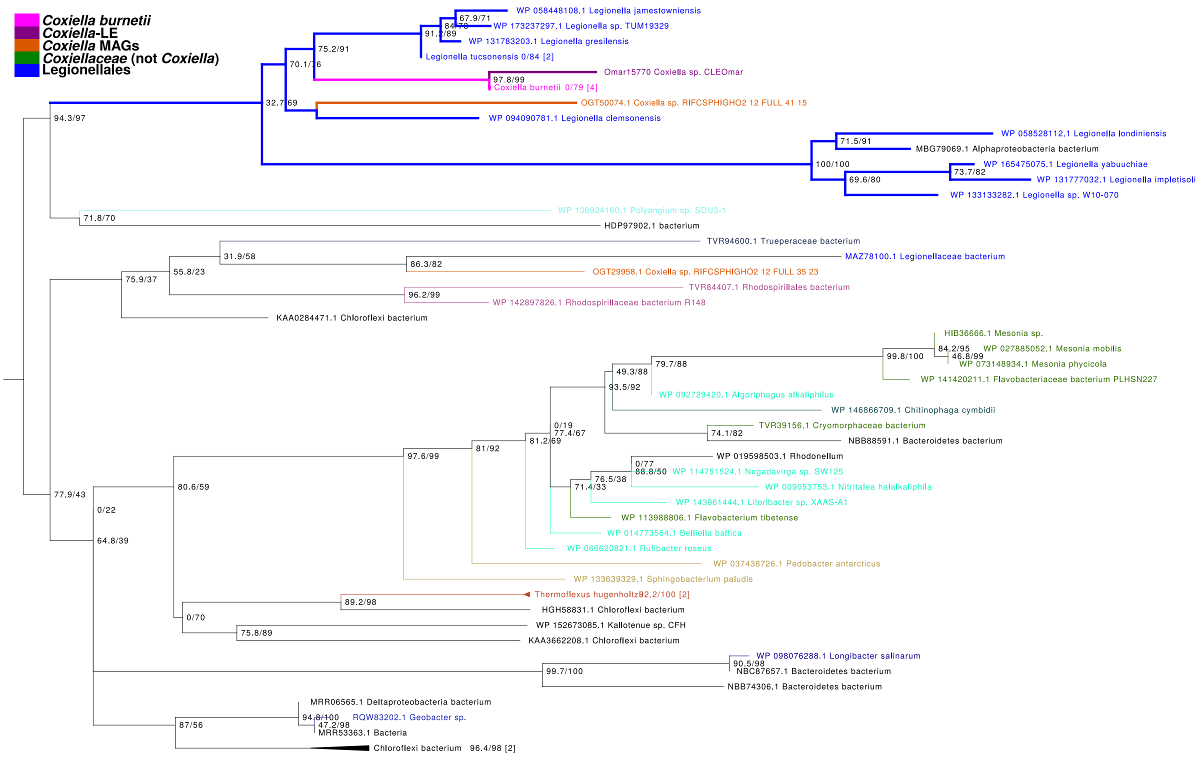


**Figure S13.** Mid-point rooted Maximum Likelihood phylogenetic tree of ShaF. Obtained ShaF Cluster of Orthologous Proteins (COPS) together with the first 50 BLASTP hits (non-redundant database, excluding *Coxiella* species) were included in the reconstruction. The clade containing ShaF from *Coxiella burnetii* is denoted by thicker lines. The numbers at each node represent 1,000 SH-aLRT (left) and ultrafast bootstrap (right) support.

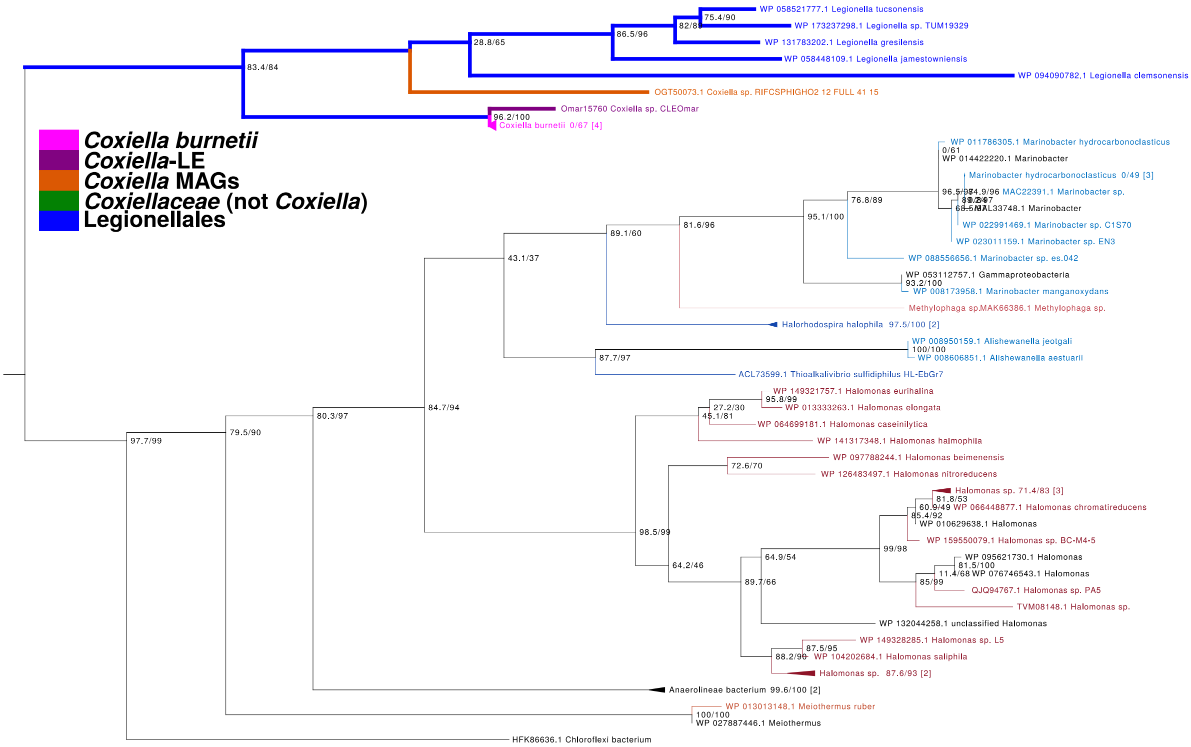


**Figure S14.** Mid-point rooted Maximum Likelihood phylogenetic tree of ShaG. Obtained ShaG Cluster of Orthologous Proteins (COPS) together with the first 50 BLASTP hits (non-redundant database, excluding *Coxiella* species) were included in the reconstruction. The clade containing ShaG from *Coxiella burnetii* is denoted by thicker lines. The numbers at each node represent 1,000 SH-aLRT (left) and ultrafast bootstrap (right) support.

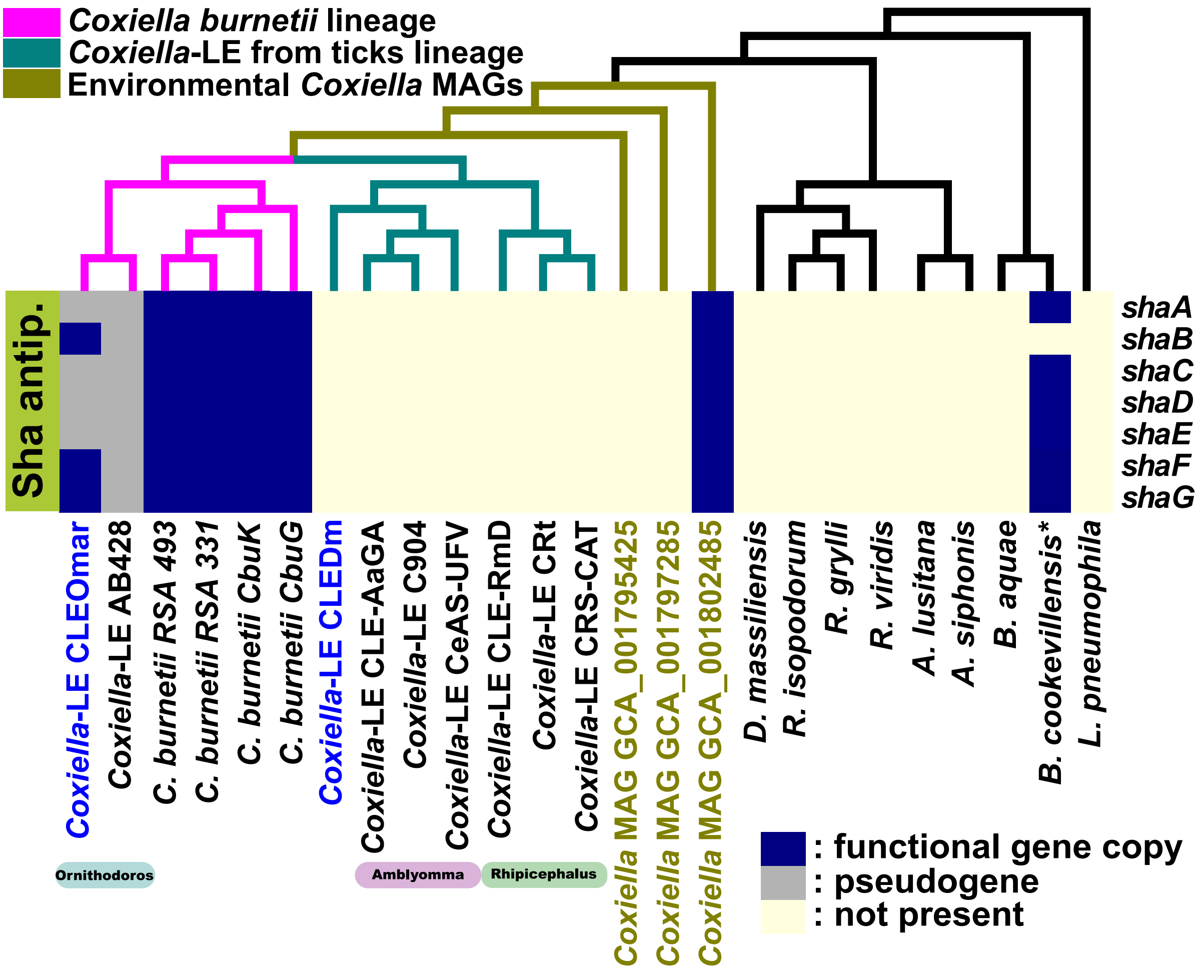


**Figure S15.** Sha/Mrp antiporter presence in selected Coxiellaceae species. *Coxiella*-LEs genomes obtained in this work are highlighted in blue. Environmental *Coxiella* MAGs basal to *C. burnetii* and *Coxiella*-LEs clade are highlighted in green. The cladogram on the top represents the phylogenetic relationships of Coxiellaceae species based on Fig 1. Species color coding is as in Fig 1. *: in *Berkiella cookevillensis*, *shA* and *shaB* are fused (*shaA'*)

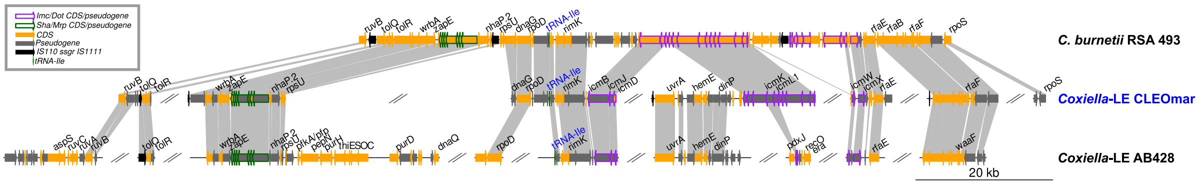


**Figure S16.** Dot/Icm T4SS genomic region from *C. burnetii* RSA 493 compared to *Coxiella*-LEs CLEOmar from *Ornithodoros maritimus* and AB428 from *O. amblus*. For draft genomes, only contigs, or regions (denoted as dotted lines) containing Dot/Icm or *Sha* genes are displayed. Gray lines connect orthologous genes. Twisted lines indicate inversions. Genes annotated as hypothetical proteins or without official names are not displayed.

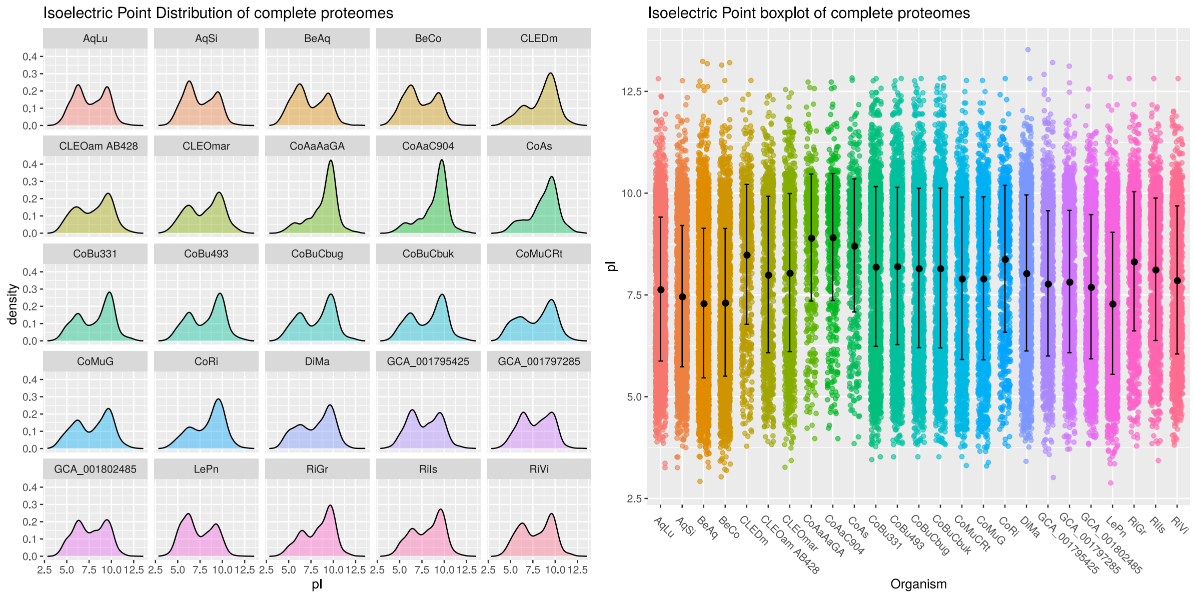


**Figure S17.** Isoelectric Point density (left) and boxplot (right) plots of full Coxiellacea proteomes. Species abbreviations: *Aquicella lusitana* SGT-39 (AqLu), *A. siphonis* SGT-108 (AqSi); *Berkiella cookevillensis* (BeCo); *B. aquae* HT99 (BeAq); *Coxiella*-LEs from *Dermacentor marginatus* CLEDm, *Ornithodoros amblus* (CLEOam AB428), *O. maritimus* (CLEOmar); *Amblyomma americanum* CLE-AaGA and C904, *A. sculptum* CeAS-UFV, *Rhipicephalus microplus* CLE-RmD, *R. turanicus* CRt, and *R. sanguineus* CRS-CAT; *C. burnetii* strains RSA 331, RSA 493, CbuK , and CbuG; environmental *Coxiella* MAGs GCA_001795425, GCA_001797285, and GCA_001802485; *Diplorickettsia massiliensis* 20B (DiMa); *Rickettsiella grylli* TrM1 (RiGr), *R. isopodorum* RCFS (RiIs), and *R. viridis* ApRA04 (RiVi); *Legionella pneumophila* Philadelphia 1 (LePn).
